# A comprehensive fitness landscape model reveals the evolutionary history and future evolvability of eukaryotic *cis*-regulatory DNA sequences

**DOI:** 10.1101/2021.02.17.430503

**Authors:** Eeshit Dhaval Vaishnav, Carl G. de Boer, Moran Yassour, Jennifer Molinet, Lin Fan, Xian Adiconis, Dawn A. Thompson, Francisco A. Cubillos, Joshua Z. Levin, Aviv Regev

## Abstract

Mutations in non-coding *cis*-regulatory DNA sequences can alter gene expression, organismal phenotype, and fitness. Fitness landscapes, which map DNA sequence to organismal fitness, are a long-standing goal in biology, but have remained elusive because it is challenging to generalize accurately to the vast space of possible sequences using models built on measurements from a limited number of endogenous regulatory sequences. Here, we construct a sequence-to-expression model for such a landscape and use it to decipher principles of *cis*-regulatory evolution. Using tens of millions of randomly sampled promoter DNA sequences and their measured expression levels in the yeast *Sacccharomyces cerevisiae*, we construct a deep transformer neural network model that generalizes with exceptional accuracy, and enables sequence design for gene expression engineering. Using our model, we predict and experimentally validate expression divergence under random genetic drift and strong selection weak mutation regimes, show that conflicting expression objectives in different environments constrain expression adaptation, and find that stabilizing selection on gene expression leads to the moderation of regulatory complexity. We present an approach for detecting selective constraint on gene expression using our model and natural sequence variation, and validate it using observed *cis*-regulatory diversity across 1,011 yeast strains, cross-species RNA-seq from three different clades, and measured expression-to-fitness curves. Finally, we develop a characterization of regulatory evolvability, use it to visualize fitness landscapes in two dimensions, discover evolvability archetypes, quantify the mutational robustness of individual sequences and highlight the mutational robustness of extant natural regulatory sequence populations. Our work provides a general framework that addresses key questions in the evolution of *cis*-regulatory sequences.

## Introduction

Changes in *cis*-regulatory elements (CREs) play a major role in the evolution of gene expression(*1, 2*). Mutations in CREs can affect their interactions with transcription factors (TFs), change the timing, location and level of gene expression, and impact organismal phenotype and fitness(*3–6*). While TFs evolve slowly because they each regulate many target genes, CREs evolve much faster and are thought to drive substantial phenotypic variation(*7–10*). Thus, understanding how *cis*-regulatory sequence variation affects gene expression, phenotype and organismal fitness is fundamental to our understanding of regulatory evolution(*6*).

A fitness function maps genotypes (which vary through mutations) to their corresponding organismal fitness values (where selection operates)(*11, 12*). A complete fitness landscape(*13–17*) is defined by a fitness function that can accurately map each sequence in a sequence space to its associated organismal fitness, ideally coupled with an approach for visualizing the complete sequence space. Partial fitness landscapes have been characterized empirically (*18–20*), often using maximum growth rate as a proxy for fitness(*l8, 21–23*). Many recent studies have favored molecular activities as fitness proxies, which are less susceptible to experimental biases and measurement noise(*24, 25*). For example, studies have now described empirical fitness landscapes of proteins(*26, 27*), adeno-associated viruses(*28*), catalytic RNAs(*29*), promoters(*30, 31*) and TF binding sites(*32, 33*), each using their respective molecular activities as indicators of fitness. In particular, the molecular activity of a promoter sequence as reflected in the expression of the regulated gene has been used to build a ‘promoter fitness landscape’(*30*). However, despite advances in high-throughput measurements, empirical fitness landscape studies remain limited to a tiny subset of the complete sequence space whose size grows exponentially with the sequence length (4^*L*^ for DNA or RNA, where *L* is the length of sequence)(*18, 30*), and often sample sequences in the local neighborhood of natural ones(*18, 19, 34*).

Predicting expression phenotype and fitness from sequence would allow us to answer fundamental questions(*34*) in evolution and gene regulation in addition to providing an invaluable bioengineering tool(*34–39*). A model relating sequence to expression comprehensively and accurately could predict how *cis*-regulatory mutations affect expression and fitness (when coupled with expression-to-fitness curves(*22, 40–42*)), design new sequences with desired characteristics, determine how quickly selection can act to reach a new expression optimum, identify signatures of the selective pressures that have shaped natural *cis*-regulatory sequences observed in extant species, visualize fitness landscapes in sequence space and characterize mutational robustness and evolvability of *cis*-regulatory sequences(*6, 18, 19, 34, 43–45*).

Here, we tackle these long-standing questions by developing a framework for studying *cis*-regulatory evolution (**Fig. 1a**) based on a *Saccharomyces cerevisiae* promoter sequence-to-expression model. We learned this model from the measured expression levels associated with tens of millions of random sequences using a deep transformer neural network. The model has exceptional predictive accuracy, which we leverage for model-guided sequence design with an evolutionary algorithm, yielding sequences that drive expression beyond the natural range in yeast. We predict (and validate experimentally) the impact of random genetic drift and the strong-selection weak-mutation regime on gene expression, show that optimizing conflicting expression objectives constrains expression adaptation even though a single expression objective can be reached with few mutations, and find that stabilizing selection on expression results in a moderation of regulatory complexity. We use the model-predicted expression differences caused by natural genetic variation in promoters to detect signatures of stabilizing selection on expression directly from regulatory sequences (without concomitant expression measurements), analogous to d_N_/d_S_ in proteins, allowing us to predict whether a gene’s expression is conserved across species and how it affects organismal fitness. Finally, we quantify the evolvability of regulatory sequences by the extent of expression changes available to each sequence by mutation. We relate sequences by their evolvability in a two-dimensional representation, distinguishing mutationally plastic and robust sequence archetypes. This representation of evolvability allows us to detect selective constraint on gene expression from individual sequences, shows that promoters of genes under stabilizing selection on gene expression tend to be mutationally robust, and provides the basis for a systematic exploration of *cis*-regulatory fitness landscapes.

**Fig. 1.**
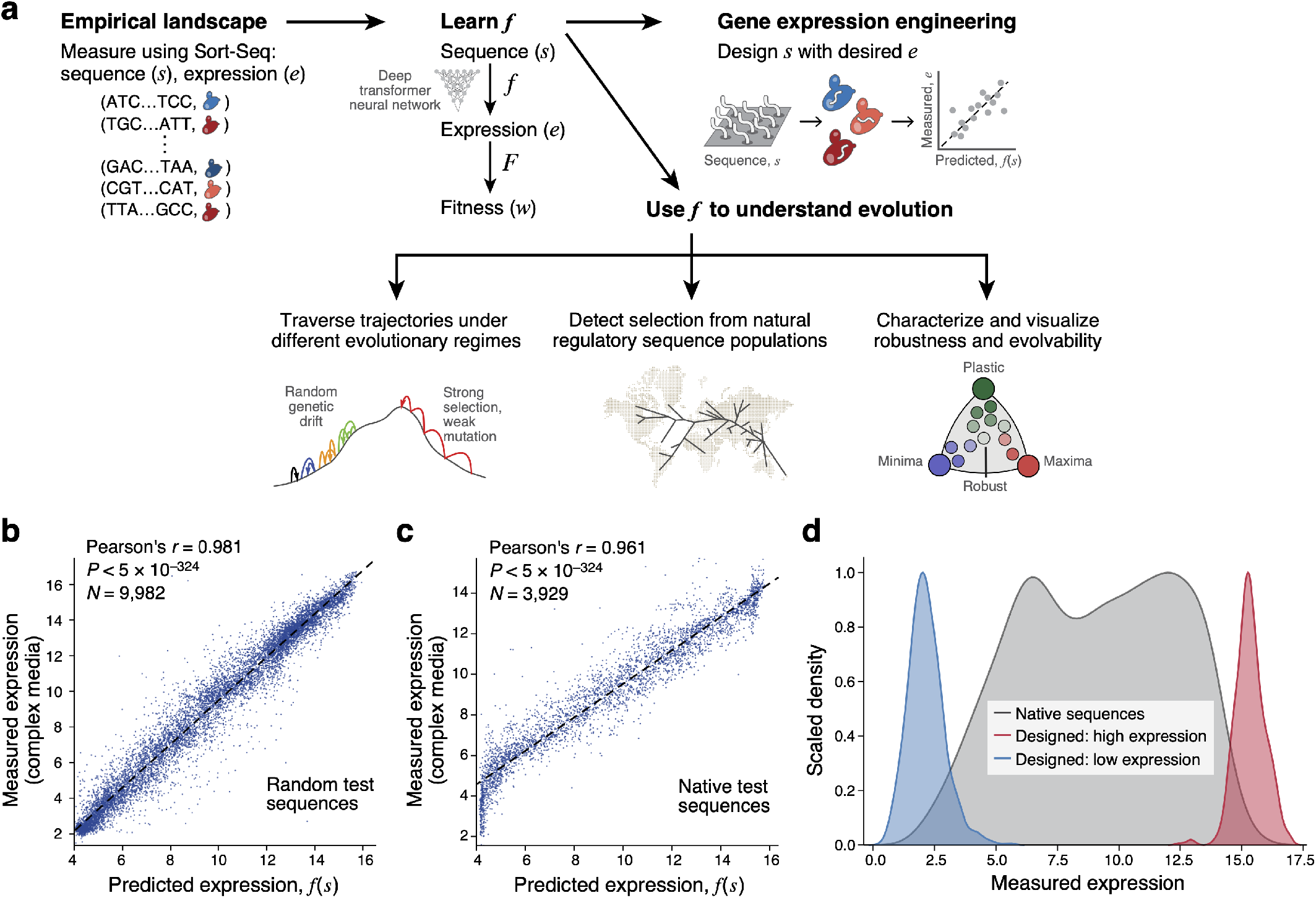
An accurate, comprehensive sequence-to-expression model enables gene expression engineering. **a,** Approach overview. **b,c,** Accurate prediction of expression from sequence. Predicted (*x* axis) and experimentally measured (*y* axis) expression in complex media for **(b)** random test sequences (sampled separately from and not overlapping with the training data) and **(c)** native yeast promoter sequences. Pearson’s *r* and associated P-values are shown. **d,** Engineering extreme expression values beyond the range of native sequences using a genetic algorithm (GA) and the sequence-to-expression model. Normalized kernel density estimates of the distributions of measured expression levels for native yeast promoter sequences (grey), and sequences designed (by the GA) to have high (red) or low (blue) expression.

## Results

### Learning a sequence-to-expression model from tens of millions of random sequences

We begin by building a model that takes DNA sequence as input and predicts expression. Here, we consider the sequence space comprising any 80 bp DNA sequence that occupies the −160 to −80 region (with respect to the Transcription Start Site (TSS)) of a promoter construct in *S. cerevisiae* (**Methods**), a critical location for TF binding(*46*) and determinant of promoter activity(*47*). To avoid biases(*19*) towards extant sequences, we measured the expression for each of over 20 million randomly sampled 80 bp DNA sequences using our previously described approach(*47*) (**Methods**). Here, we clone random sequences into a YFP expression vector, transform them into yeast grown in a defined medium (SD-Ura, synthetic defined lacking uracil; **Methods**), sort the yeast into 18 expression bins, and sequence the promoters in the yeast in each bin to estimate expression.

We learned the model using a deep transformer neural network that can predict expression values from sequence, with a model architecture (**Methods**, **Supplementary Fig. S1**) designed to reflect known aspects of *cis*-regulation(*48, 49*). Briefly, the model has three blocks, each consisting of multiple layers, and analogous to different biological aspects. The first is a convolutional block with three layers, which identifies sites that are important for computing the expression target, and are analogous to a TF scanning the length of the sequence for binding sites. The first layer learns an abstract representation of first-order TF-sequence interactions by operating with convolutional kernels on the sequence, scanning the forward and reverse strands separately to generate strand-specific features (each individual kernel in the first layer can be thought of as learning the motif of one TF, or a combined representation of the motifs)(*50–53*); the second can capture interactions between strands, by using a 2D convolution on the combined features from the individual strands; and the third layer can capture higher order interactions, such as TF-TF cooperativity. The second block is analogous to combining the biochemical activities of multiple bound TFs and accounting for their positional activities. It first uses a transformer-encoder with a multi-head self-attention module(*54*) to capture relations between features extracted by the convolutional block at different positions in the sequence, by attending to them simultaneously using a scaled dot product attention function. Here, the model can learn ‘where to look’ within the sequence. Then, a bidirectional Long Short-Term Memory (LSTM) layer in this block captures long range interactions between the sequence regions. Finally, a multi-layer perceptron block can capture cellular operations that occur after TFs are recruited to the promoter sequence, by pooling all the features extracted from the sequence through the previous layers and learning a scaling function that transforms these abstract feature representations of biomolecular interactions into an expression estimate.

### The model generalizes in the sequence space to accurately predict expression from sequence

To show that the learned model can generalize, we predicted the expression of new test sequences not seen during model training, and compared them to their experimentally measured levels (**Methods**, assayed in the same SD-Ura defined media used for generating the training data). We obtained exceptionally accurate predictions both when testing on a 5,351 random sequence test set (Pearson’s *r* = 0.969, *P* < 5*10^-324^, **Supplementary Fig. S2a**) and when testing on the 3,978 native yeast promoter sequences (−160 bp to −80 bp relative to the TSS of native yeast genes) for which we quantified expression using our assay (Pearson’s *r* = 0.95, P < 5*10^-324^, **Supplementary Fig. S2b**).

As a contrasting growth condition, we trained a second model using another 30 million sequence-expression pairs measured separately in a complex growth medium (YPD, **Methods**). Here, too, we observed excellent performance on 9,982 random test sequences (Pearson’s *r* = 0.981, *P* < 5*10^-324^, **Fig. 1b**) or 3,929 native yeast promoter sequences (Pearson’s *r* = 0.961, *P* < 5*10^-324^, **Fig. 1c).** These results represent a decrease in error of 33%-50% compared to the performance of our biochemical models(*47*) when learned here from the same data, highlighting the superior predictive power of the deep transformer model. Moreover, the expression measurements were highly correlated for the same sequences between the two media (Pearson’s *r* = 0.978, **Supplementary Fig. S3a**) and the model trained on the defined medium predicted expression in the complex medium well (Pearson’s *r* = 0.970, **Supplementary Fig. S3b**). However, for some sequences we expect differences between growth conditions, as we study below.

### Model-guided expression engineering beyond the range of native expression

We leveraged the high predictive accuracy of the model for a synthetic biology application of gene expression engineering, by using our model as the ‘fitness function’ for a genetic algorithm (GA) to design sequences with extreme expression values. We initialized the GA with a population of 100,000 randomly-generated samples from the sequence space, and simulated 10 generations to maximize (or minimize) the expression output (**Methods**). To test whether the designed sequences from the GA indeed achieve their predicted extreme expression levels, we synthesized the 500 sequences with the top predicted maximum (or minimum) expression levels and tested them experimentally. The GA-designed sequences drove, on average, more extreme expression than >99% of native sequences (99.6% for high expressing; 99.3% for low), with ~20% of designed sequences more extreme than any native sequence tested (23.5% for high; 18.4% for low) (**Fig. 1d**). Thus, our sequence-to-expression model can be used for gene expression engineering.

### Expression divergence under random genetic drift

We next assessed the impact on expression of different evolutionary scenarios: random drift, stabilizing selection, and directional selection for extreme expression levels, as well as for two opposing expression requirements (**Fig. 2**). In each case we simulated the scenario using our model to predict the expression phenotype of each sequence, and then tested the model’s evolved sequences experimentally, where possible.

**Fig. 2.**
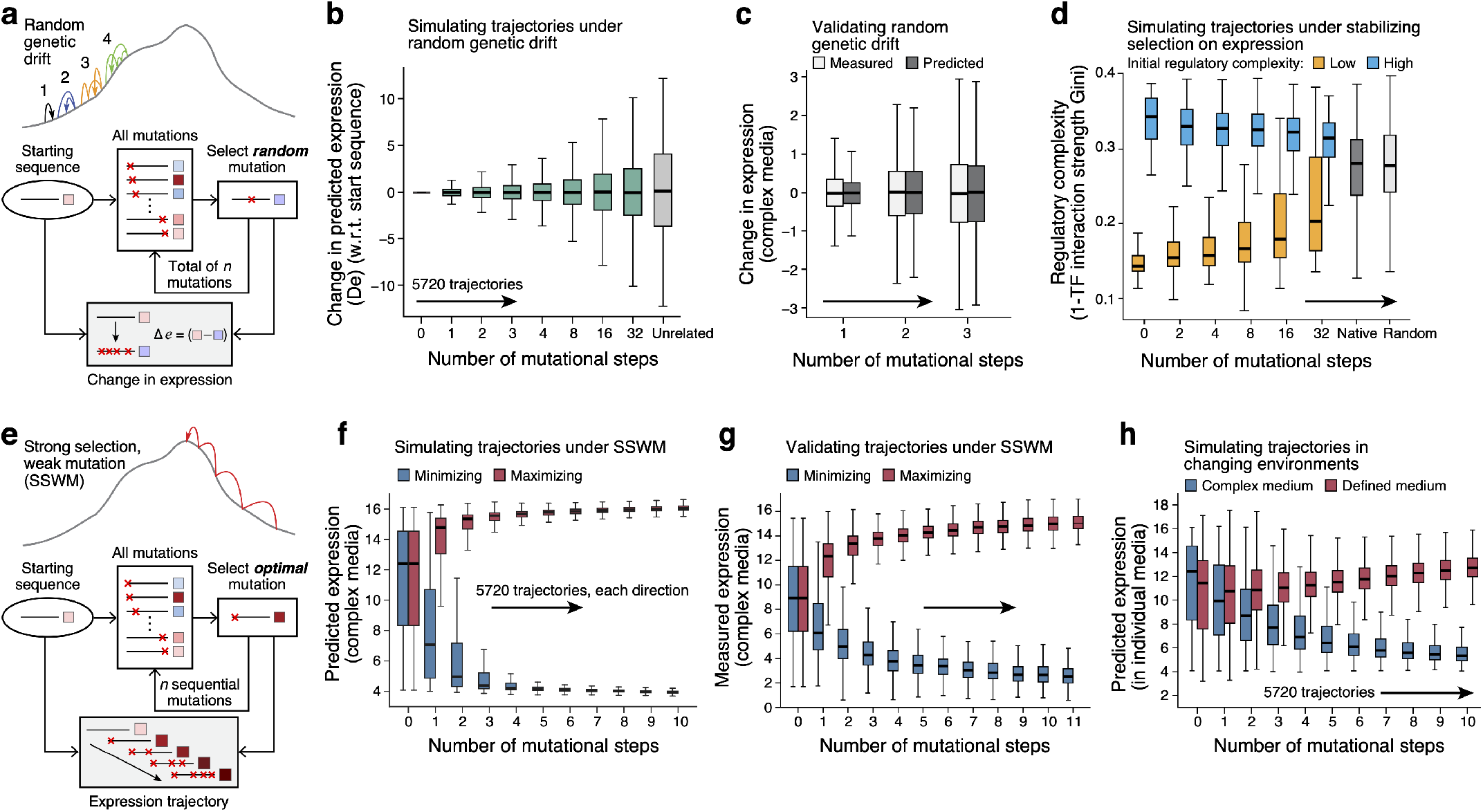
Characterizing the effects of random drift, stabilizing and directional selection on *cis*-regulatory sequences with the sequence-to-expression model. **a-c,** Expression divergence under random genetic drift. **a.** Simulating trajectories. Top: An imaginary fitness landscape with trajectories for one (black), two (blue), three (orange), and four (green) random mutations. Bottom: Simulation procedure. **b,** Predicted expression divergence under random genetic drift. Distribution of the change in predicted expression (*y* axis) for 5,720 starting sequences at each mutational step (*x* axis) for trajectories simulated under random mutational drift. Silver bar: differences in expression between unrelated sequences. Midline: median; boxes: interquartile range; whiskers: 1.5x interquartile range. **c,** Experimental validation. Distribution of measured (light grey) and predicted (dark gray) changes in expression in complex media (*y* axis) for the synthesized sequences at each mutational step (*x* axis) from predicted mutational trajectories under random mutational drift. Midline: median; boxes: interquartile range; whiskers: 1.5x interquartile range. **d,** Stabilizing selection on gene expression leads to moderation of regulatory complexity extremes. Regulatory complexity (*y* axis) for sequences from sequential mutational steps (*x* axis) under stabilizing selection to maintain the starting expression levels, where the regulatory interactions of starting sequences are initially complex (blue; n=47) or simple (orange, n=64), in complex media (YPD). Right bars: regulatory complexity for native (dark gray) and random (light gray) sequences. Midline: median; boxes: interquartile range; whiskers: 1.5x interquartile range. **e-g,** Sequences under SSWM can rapidly evolve to an expression optimum. **e.** Simulating trajectories under SSWM. Top: An imaginary fitness landscape with one trajectory to achieve an expression optimum. Bottom: Simulation procedure. **f,** Predicted expression evolution under SSWM. Distribution of predicted expression levels (*y* axis) in complex media at each mutational step (*x* axis) for sequence trajectories under SSWM favoring high (red) or low (blue) expression, starting with 5,720 native promoter sequences. Midline: median; boxes: interquartile range; whiskers: 1.5x interquartile range. **g,** Experimental validation. Experimentally measured expression distribution in complex media (*y* axis) for the synthesized sequences at each mutational step (*x* axis) from predicted mutational trajectories under SSWM, favoring high (red) or low (blue) expression. Midline: median; boxes: interquartile range; whiskers: 1.5x interquartile range. **h,** Competing expression objectives constrain expression adaptation. Distribution of predicted expression (*y* axis) in complex (blue) and defined (red) media at each mutational step (*x* axis) for a starting set of 5,720 native promoter sequences optimizing for high expression in defined media (red) and simultaneous low expression in complex media (blue).

We first simulated random drift of regulatory sequences, with no selection on expression levels. We randomly introduced a single mutation in each starting sequence, repeated this process for multiple consecutive generations, and then used our model to predict the difference in expression between the mutated sequences in each trajectory relative to the corresponding starting sequence (**Fig. 2a-c**).

Expression levels diverged as the number of mutations increased, with 32 mutations resulting in nearly as different expression from the original sequence as two unrelated sequences (**Fig. 2b**). We validated our results experimentally by synthesizing sequences with zero to three random mutations and measuring their expression in our assay (**Methods**). The experimental measurements closely matched our predictions in both complex (**Fig. 2c**) and defined (**Supplementary Fig. S2c**) media, with excellent agreement in both expression change (Pearson’s *r*: 0.877 and 0.849, respectively; **Supplementary Fig. S2f,g**) and expression level (Pearson’s *r*: 0.974 and 0.963 respectively; **Extended Fig. 2j,k**).

### Stabilizing selection on expression leads to a moderation of regulatory complexity extremes

Although gene regulatory networks often appear to be highly interconnected(*47, 55–57*), the sources of this regulatory complexity and how it changes with the turnover of regulatory mechanisms(*58*) remain unclear. We thus used our model to study the evolution of regulatory complexity in the context of stabilizing selection, which favors the maintenance of existing expression levels. First, we used an interpretable biochemical model we previously developed(*47*) to quantify regulatory complexity, defined as 1 minus the Gini coefficient of TF regulatory interaction strengths (**Methods**). Starting with native sequences whose regulatory complexity is either extremely high (many TFs with similar contributions to expression) or low (few TFs contribute disproportionately to expression), we simulated how regulatory complexity changed under stabilizing selection on gene expression (with each starting set of sequences chosen to span a range of expression levels). We introduced single mutations into each starting native sequence for each of 32 consecutive generations, identified the sequences that conserved the original expression level and selected one of them at random for the next generation. To ensure that expression levels remained unchanged, we experimentally measured expression for generations 2, 4, 8, 16, and 32, and excluded any trajectories for which any of these differed from the original expression level by more than 1. We then asses the regulatory complexity of the evolved sequences as before.

We found that as random mutations accumulated, the regulatory complexity of sequences starting at both complexity extremes shifted towards moderate regulatory complexities (**Fig. 2d**, rightmost blue and orange), closer to the averages for both random and native sequences (**Fig. 2d,** greys). This suggests that stabilizing selection on expression leads to a moderation of regulatory complexity, resulting from gradual drift in the roles of the different regulators. Further, the overall distribution of regulatory complexity of native yeast promoters is similar to that of random sequences (**Fig. 2d,** grey boxes), suggesting that the regulatory complexity of native sequences primarily reflects their sampling from the space of sequences with equivalent expression outcomes.

### Directional selection for extreme expression requires few mutations

To study the impact of directional selection on gene expression, we used our model to simulate the strong-selection weak-mutation (SSWM) regime(*59, 60*) (**Fig. 2e, Methods**), where each mutation is either beneficial or deleterious (strong selection, with mutations surviving drift and fixing in an asexual population), and mutation rates are low enough to only consider single base substitutions during adaptive walks(*61, 62*) (weak mutation). Briefly, starting with the set of all native sequences, at each iteration (generation), for a given starting sequence of length *L*, we consider all of its 3*L* single base mutational neighbors, use our model to assess their expression, and take the sequence with the largest increase (or separately, decrease) in expression at each iteration (generation) as the starting set of sequences for the next generation (**Fig. 2e, Methods**).

Sequences that started with diverse initial expression levels rapidly evolved to high (or separately, low) expression, with the vast majority evolving close to saturating extreme expression levels within 3-4 mutations in both the complex (**Fig. 2f**) and defined (**Supplementary Fig. S2d**) media models. We validated these trajectories experimentally for select series of sequences (**Fig. 2g, Supplementary Fig. S2e**), measuring the expression driven by synthesized sequences from several generations along simulated mutational trajectories for complex media (10,322 sequences from 877 trajectories) and defined media (6,304 sequences from 637 trajectories). We again observed extreme expression within 3-4 mutational steps, with high agreement between measured and predicted expression change (**Supplementary Fig. S2h,i**; Pearson’s *r*: 0.973 and 0.956, respectively) and expression levels (**Supplementary Fig. S2l,m**; Pearson’s *r*: 0.981 and 0.974) along the trajectories in both the complex and defined media.

### Conflicting expression objectives in different environments constrain expression adaptation

In contrast to the rapid evolution towards expression extremes, evolution to satisfy two opposing expression requirements (one in each growth media) is more constrained. A concrete example is the expression of the *URA3* gene, which codes for an enzyme in the uracil synthesis pathway: in defined media lacking uracil organismal fitness *increases* with increased expression, but in complex media containing 5-FOA fitness *decreases* with increased expression, due to Ura3-mediated conversion of 5-FOA to toxic 5-fluorouracil (**Supplementary Fig. S3c**). To study this regime, we used our model with the set of all native sequences (and separately, a set of random sequences) as starting sequences and simulated trajectories under the SSWM regime, simultaneously optimizing for two competing objectives: maximize expression in the defined medium, while minimizing it in the complex medium. Here, we maximize the difference in expression between the two conditions at each iteration, assuming that the cells are exposed to both environments before the mutations can reach fixation (**Methods**). While the difference in expression increased with each generation (**Supplementary Fig. S3d,e**), the vast majority of sequences achieved neither the maximal nor the minimal expression in either condition (**Fig. 2h**, **Supplementary Fig. S3f**), for both native and random starting sequences. Interestingly, after 10 generations, the evolved sequences became enriched for motifs for TFs involved in nutrient sensing and metabolism, compared to the starting sequences (**Supplementary Fig. S3g**). Thus, while evolving a sequence to achieve a single expression optimum requires very few mutations, encoding multiple opposite objectives within the same sequence is more difficult, suggesting that conflicting expression objectives in different environments constrain expression adaptation.

### The Expression Conservation Coefficient (ECC) detects signatures of stabilizing selection on gene expression using natural genetic variation in *cis*-regulatory DNA

We next applied our sequence-to-expression model to detect evidence of selective pressures on natural regulatory sequences, inspired by the way in which the ratio of non-synonymous (“non-neutral”) to synonymous (“neutral”) substitutions (d_N_/d_S_) in protein coding sequences is used estimate the strength and mode of natural selection(*63*). By analogy(*64–66*), for regulatory sequences(*6*), we used our model to assess the impact (a continuum between “nonneutral” and “neutral”) of naturally occurring regulatory mutations on gene expression, compared to that expected with random mutations, and summarize this with an Expression Conservation Coefficient (ECC) (**Methods**). To compute the ECC, we compared, for each gene’s promoter, the standard deviation of the expression distribution predicted by the model for a set of naturally varying orthologous promoters (σ_B_) to the standard deviation of the expression distribution predicted for a matched set of random variation introduced to that promoter (σ_C_; **Fig. 3a**). The random variation was generated by placing random mutations within the gene’s promoter consensus (the most abundant base at each position in the orthologous set), while preserving the Hamming distance distribution observed in the natural sequences (**Methods**). We define the ECC for a gene as log(σ_c_/σ_B_), such that a positive ECC indicates stabilizing selection on expression (lower variance in the native sequences), a negative ECC indicates diversifying (disruptive) or directional (positive) selection, and values near 0 suggest neutral drift (see **Methods** for definitions).

**Fig. 3.**
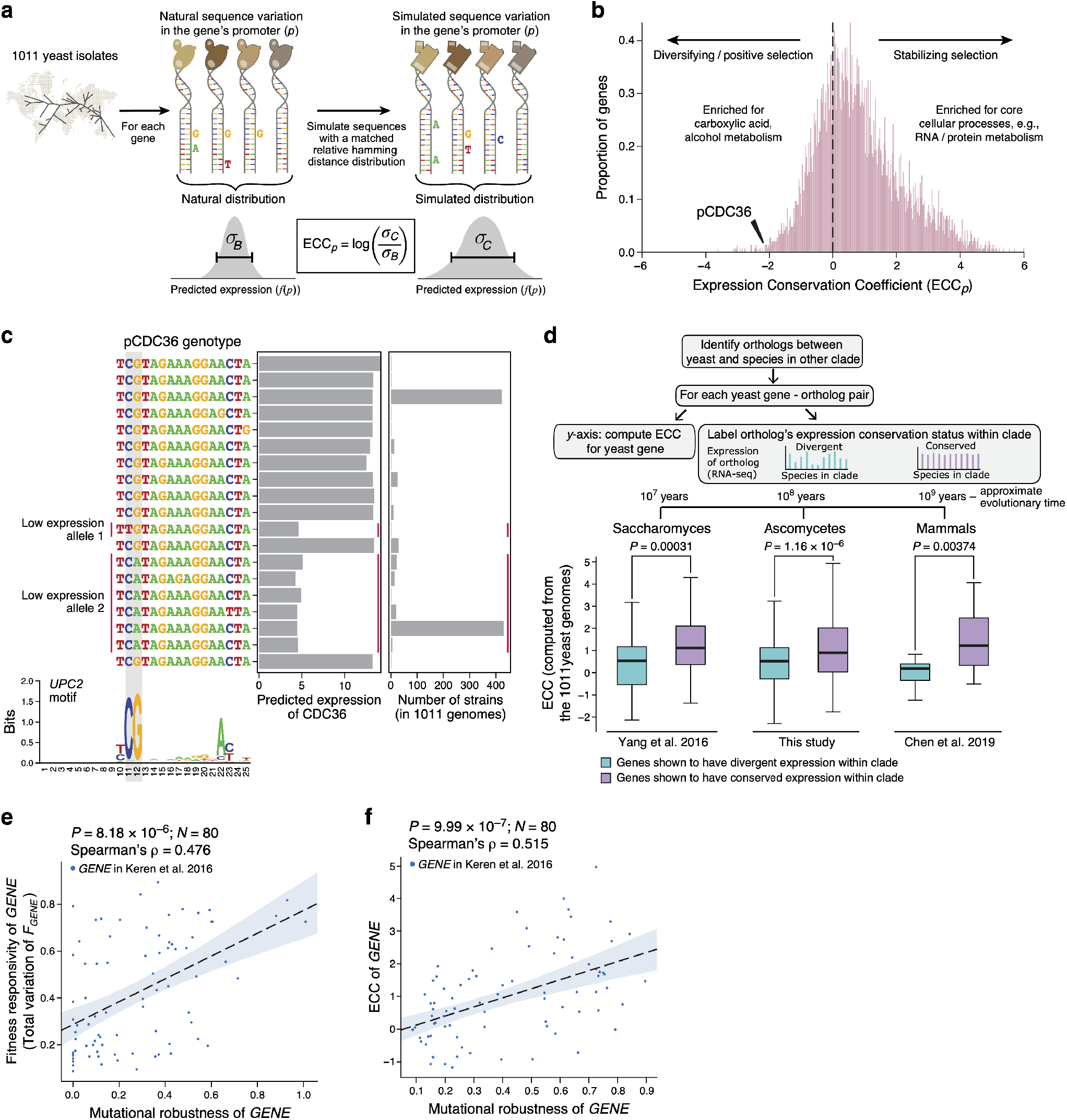
The Expression Conservation Coefficient (ECC) detects signatures of stabilizing selection on gene expression using natural genetic variation in *cis*-regulatory DNA. **a-c, a,** ECC calculation from 1,011 *S. cerevisiae* genomes^60^. **b,** ECC distribution for *S. cerevisiae* genes. Frequency distribution of ECC values (*x* axis). Dashed line distinguishes regions corresponding to disruptive/positive selection (left) and stabilizing selection (right) and GO terms enriched by the ECC ranking. Arrowhead: ECC value for the CDC36 promoter sequence. **c,** Convergent regulatory evolution in the CDC36 promoter. Predicted expression (*x* axis, left bar plot) and associated number of strains (*x* axis, right bar plot) of all alleles among the analyzed CDC36 promoter sequence within 1,011 yeast isolates, along with an alignment of their UPC2 binding site sequences (left; UPC2 binding motif below). Red vertical lines: two independently evolved low-expressing alleles. Grey vertical boxes: key positions in the UPC2 motif with single nucleotide polymorphisms. **d**, Distribution of ECC (*y* axis, calculated from 1,011 S. cerevisiae genomes, top left) for *S. cerevisiae* genes whose orthologs have divergent (blue) or conserved (purple) expression (within *Saccharomyces* (left), Ascomycota (middle), or mammals (right) (as determined by cross species RNA-seq, top right). P-values: two-sided Wilcoxon rank-sum test. Midline: median; boxes: interquartile range; whiskers: 1.5x interquartile range. **e,f**, Genes whose expression changes have stronger effects on organismal fitness have mutationally robust regulatory sequences. Mutational robustness (*x* axes) and fitness responsivity (**e**, *y* axis) or ECC (**f**; *y* axis) for each of 80 genes (points) for which the expression-to-fitness curves were quantified^21^. Spearman’s *ρ* and associated P-values are shown.

We calculated the ECC for 5,569 *S. cerevisiae* genes using the natural variation observed in the −160 to −80 regions of over 4.73 million orthologous promoter sequences from the 1,011 *S. cerevisiae* isolates(*67*) (**Fig. 3a,b, Supplementary Table 1**). Over 70% of promoters had positive ECCs, suggesting stabilizing selection (and conserved expression) (binomial test P < 10^-215^) (**Fig. 3b**), consistent with previous reports based on direct measurements of gene expression(*68, 69*). Genes with high ECCs were enriched in highly-conserved core cellular processes (*e.g*., RNA and protein metabolism(*70*)) (**Fig. 3b, Supplementary Table 2**), and those with low ECCs were most enriched in processes related to carboxylic acid and alcohol metabolism (**Fig. 3b, Supplementary Table 2**), potentially reflecting adaptation of fermentation genes to the diverse natural and industrial settings from which these isolates were collected(*67*).

A striking example of predicted positive selection is the promoter of CDC36 (ECC= −2.138, **Fig. 3b**), which has common natural alleles with either low or high (predicted) expression across the isolates (**Fig. 3c**). Analysis of CDC36 promoter sequences (**Methods**) suggests that low-expression evolved at least twice independently, resulting in two distinct variants with reduced expression (**Fig. 3c, allele 1 and 2**). Interrogation of our previously published biochemical model to identify factors impacting these expression differences (**Supplementary Fig. S4a**) suggested that both low-expression alleles are explained by disruption of the same binding site for Upc2p, an ergosterol sensing TF (**Fig. 3c**). The two mutation events at adjacent nucleotides in one TF binding site support the hypothesis that these two independent mutations result from convergent evolution of a common new CDC36 regulatory and expression phenotype, which is captured by the low ECC of CDC36.

### The ECC is consistent with cross-species RNA-seq and expression-to-fitness measurements

ECC values were consistent with expression conservation as measured for yeast orthologs across clades at short (*Saccharomyces*), medium (Ascomycota), or long (mammals) evolutionary scales (**Fig. 3d**). In *Saccharomyces*, orthologs with conserved expression levels across *Saccharomyces* species (as measured by RNA-seq(*71*)) had significantly higher ECC (computed from the 1,011 yeast isolates) than genes whose expression was not conserved (two-sided Wilcoxon rank-sum test P = 3.1*10^-4^) (**Fig. 3d, bottom left, Methods**). Next, we performed RNA-seq across 11 Ascomycota yeast species (**Methods**), and found that orthologs with conserved expression across Ascomycota had significantly higher ECC values (**Fig. 3d, bottom center**, P = 1.16*10^-6^). Finally, genes with high ECC values in the 1,011 *S. cerevisiae* isolates also reflected expression conservation of their orthologs (one-to-one or one-to-many) within mammals(*72*) (**Fig. 3d, bottom right**, two-sided Wilcoxon rank-sum test P = 0.00374, **Methods**). Thus, while yeast-mammal orthologs are likely critical to an organism’s fitness, those under stabilizing selection for expression in yeast (by the ECC) tend also to be more conserved in expression across mammals (by RNA-seq). Thus, the ECC quantified stabilizing selection on expression in yeast and may even predict stabilizing selection on orthologs’ expression in other species.

Genes with higher ECCs also had a stronger effect on organismal fitness in *S. cerevisiae* upon changing their gene expression level, as reflected by our interrogation of previously measured expression-to-fitness relationships(*22*). Specifically, we analyzed the empirically-determined relationships between the expression levels of each of the 80 genes to organismal fitness in a published experiment(*22*), using the total variation of the expression-fitness curve as a ‘fitness responsivity’ score of how fitness depends on expression (**Supplementary Fig. S5**, **Methods**). Fitness responsivity was significantly positively correlated with the ECC (**Supplementary Fig. S4b**, Spearman ρ = 0.326, *P* = 0.003). Additionally, we find the same qualitative relationship between a gene’s ECC and fitness responsivity as reported for other genes, including *LCB2* (ECC 2.15 and high fitness responsivity(*42*)) and *MLS1* (ECC −1.32 and extremely low fitness responsivity(*41*)). Notably, fitness responsivity was not associated with regulatory sequence divergence *per se* across the promoter sequence (as estimated by the mean Hamming distance among orthologous promoters, **Methods, Supplementary Fig. S4c**, Spearman ρ = 0.083, *P* = 0.46), suggesting that while stabilizing selection on gene expression (as determined by the ECC) can shape the types of mutations that accumulate in the population, it may have little effect on the overall rate at which mutations accumulate in promoter regions within populations.

### Mutational robustness of gene promoters under stabilizing selection on expression

While a gene’s ECC (computed from the natural genetic variation in regulatory DNA) represents the imprint of its evolutionary history, its mutational robustness (assessed directly from the gene’s promoter sequence) should describe how *future* mutations would affect its expression. We defined the mutational robustness of a sequence length *L*, as the percent of its 3*L* single nucleotide mutational neighbors predicted to result in a *negligible* change in expression (**Supplementary Fig. S4d**, **Methods**), following previous descriptions of mutational robustness(*73–75*).

The mutational robustness of a gene’s promoter sequence was positively correlated with the gene’s fitness responsivity (**Fig. 3e**, Spearman ρ = 0.476, P = 8.18*10^-6^), suggesting that fitness-responsive genes have evolved more mutationally robust regulatory sequences. Mutational robustness which, unlike the ECC, is computed for single sequences without a set of variants across a population, was also correlated to the ECC (**Fig. 3f**, Spearman ρ = 0.515, P = 9.99*10^-7^). Similarly, the promoter sequences of yeast genes with conserved expression across *Saccharomyces* strains(*71*), Ascomycota species, or mammals(*72*) had higher mutational robustness (*P* = 8.4*10^-3^, 6.5*10^-5^, and 0.017, respectively, two-sided Wilcoxon rank-sum test). Thus, genes whose expression level are under stabilizing selection have regulatory sequences that are more robust to the impact of mutations (which may reflect their history and constrain their future).

### A characterization of *cis*-regulatory evolvability captures evolutionary properties of the sequence space

Mutational robustness can facilitate evolvability, the ability of a system to generate heritable phenotypic variation, by allowing the exploration of novel genotypes that may facilitate adaptation(*44, 73–75*). To characterize regulatory evolvability, we extended our description of mutational robustness to develop a general representation of the evolutionary properties of regulatory sequences. To this end, we represent each sequence as a sorted vector of expression changes (predicted by our model) that are accessible through single base substitutions to that sequence: we sort the expression changes associated with the single base changes in every possible position to obtain a monotonically increasing vector of length 3*L* for each sequence of length *L* (here, *L*=80; 3*L*=240; **Fig. 4a**, left, **Methods**). We term this representation the ‘evolvability vector’, in line with previous definitions of evolvability(*44, 73*), however its relationship with evolvability is context-dependent. Sequences for which mutations change expression (*i.e*., their evolvability vectors have a large magnitude) are evolvable in the sense that they can adapt to new expression optima easily, but under stabilizing selection the majority of mutations in such sequences would be maladaptive, limiting regulatory program evolvability. Alternatively, sequences in which mutations tend to preserve expression are less evolvable in terms of their expression level, but are more evolvable in their regulatory program since more mutations can be tolerated.

**Fig. 4.**
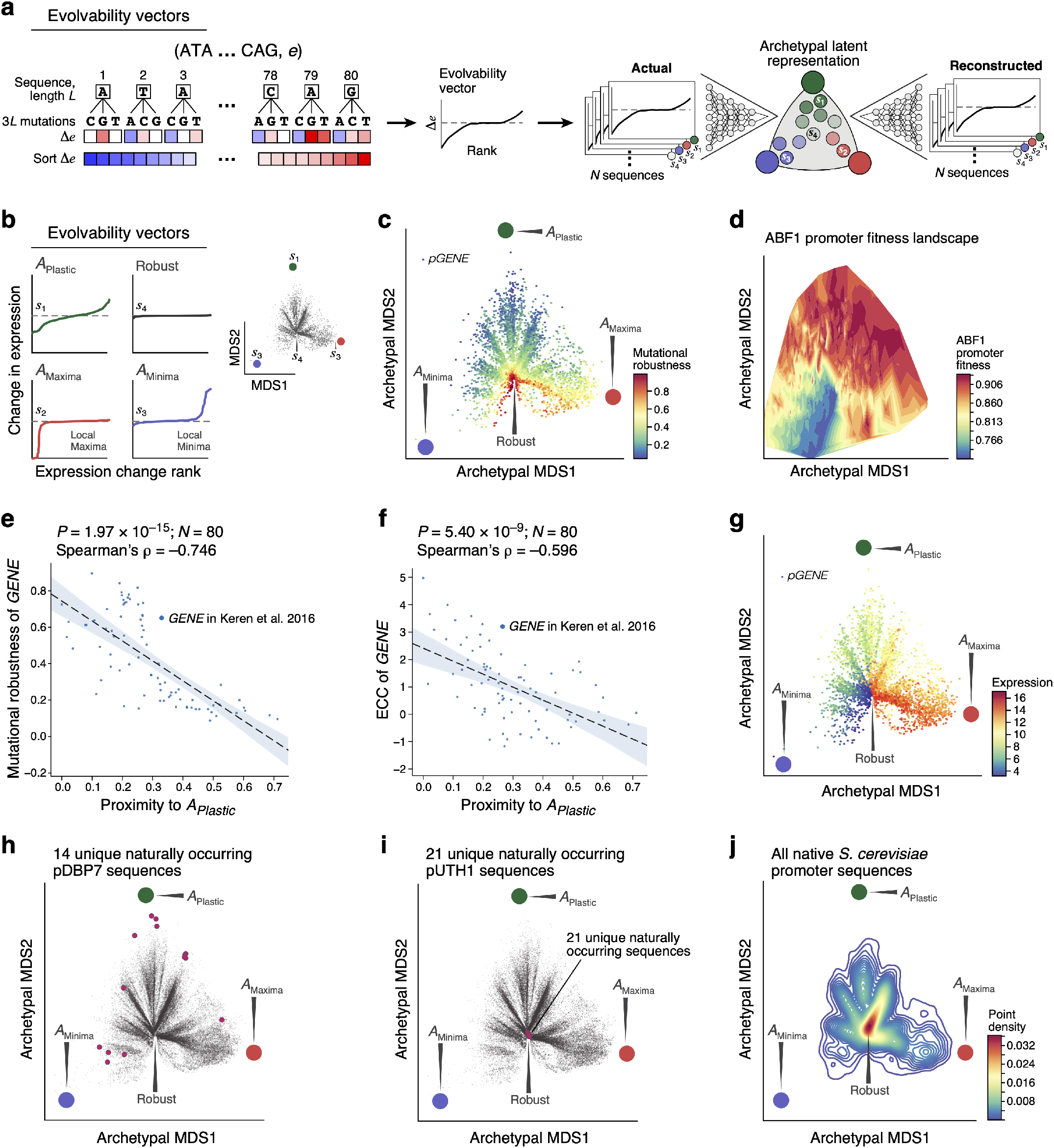
A characterization of cis-regulatory evolvability captures evolutionary properties of the sequence space and enables the visualization of fitness landscapes. **a,** Characterizing *cis*-regulatory evolvability by computing an evolvability vector and its archetypal representation. Left and middle: Generation of evolvability vectors for a given sequence. Right: training an autoencoder with evolvability vectors to generate an archetypal representation that is bounded by a simplex, and can be projected onto a 2D MDS-embedding of archetype space to visualize sequence spaces. **b,** Evolvability archetypes discovered by the autoencoder. Left: Evolvability vectors of the rank ordered (*x* axis) predicted change in expression (*y* axis) for native sequences closest to each of the plastic (green), maxima (red) or minima (blue) archetypes and the ‘robustness cleft’ (black). Right: all native yeast (*S. cerevisiae* S288C) promoter sequences (grey points) projected onto the archetype space by their evolvability vectors. Evolvability archetypes (colored circles) and their closest native sequences (s_1_-s_4_ as on left) are marked. **c,** Evolvability landscape captures mutational robustness. Evolvability vectors (points) of all native yeast promoter sequences projected onto the archetype space (coloured circles, as in **b**) and colored by mutational robustness. **d,** Visualizing the *ABF1* promoter fitness landscape. Promoter sequences represented by their respective evolvability vectors are projected onto the archetype space and colored by their associated fitness as reflected by their predicted growth rate relative to the wildtype (color, **Methods**), estimated by first mapping sequences to expression with our model and then expression to fitness as measured and estimated previously^21^. **e,f**, The evolvability vectors’ archetypal representation predicts expression conservation from solitary sequences. Proximity to the plastic archetype (A_Plastic_) (*x* axis) and mutational robustness (**e**, *y* axis) or ECC (**f**, *y* axis), for each of 80 genes with measured fitness responsivity. Top right: Spearman’s *ρ* and associated P-value. **g,** Evolvability landscape captures expression levels. Evolvability vectors (points) of all native yeast promoter sequences projected onto the archetype space (colored circles, as in b) and colored by predicted expression level. **h,i**, Plastic promoter sequences dynamically traverse the archetype space. Evolvability vector projections of native sequences (points) from all 1,011 *S. cerevisiae* isolates. Red points: natural promoter sequence variants for *DBP7*, the promoter closest to the plastic archetype (**h**) and for *UTH1*, the promoter closest to the robustness cleft (**i**). **j**, The robustness of native promoter sequences. Density (color) of all native yeast promoter sequences when their evolvability vectors are projected onto the archetype space.

We next determined whether *cis*-regulatory evolvability vectors fell into distinct classes by identifying evolvability archetypes: extreme canonical patterns of expression change in mutational neighborhoods. Using our model, we computed evolvability vectors for a new random sample of a million sequences and then embedded these evolvability vectors into a two-dimensional archetypal(*76–78*) latent space using an autoencoder(*79*) (**Fig. 4a**, right, **Methods**). This archetypal latent space is represented as a convex polyhedron whose vertices represent evolvability archetypes; each sequence can be represented as a single point within this space. This characterization of evolvability allows us to encode and visualize sequences by their evolvability in the context of a fitness landscape.

Three archetypes captured most of the variation in evolvability vectors (**Supplementary Fig. S6a,b**; **Methods**), corresponding to local expression minimum (A_Minima_), local expression maximum (A_Maxima_), and plastic expression (A_plastic_) (**Fig. 4b**). A_Minima_ and A_Maxima_ correspond to sequences where most 3*L* mutational neighbors do not change expression, and the ones that do, increase it (for A_Minima_) or decrease it (for A_Maxima_). Conversely, for the plastic archetypal sequences, most 3*L* mutational neighbors change expression and are equally likely to decrease or increase it (**Fig. 4b**). In addition to these three archetypes, mutationally robust sequences were present as a central cleft in the archetypal latent space (**Fig. 4b,c**; “Robust”). Combining our sequence-to-expression model with the expression-to-fitness curves characterized previously(*22*), and integrating them with our two-dimensional representation of evolvability, we now have a way of visualizing promoter fitness landscapes (**Fig. 4d**, **Supplementary Fig. S7**, **Methods**).

When embedding the evolvability vectors for native yeast sequences into the learned archetypal latent space, there was a strong negative correlation between a sequence’s proximity to the plastic archetype and its mutational robustness (**Fig. 4e**, **Supplementary Fig. S6c**; Spearman’s ρ = −0.746, P = 1.97*10^-15^), the ECC (**Fig. 4f**, **Supplementary Fig. S6d,e**; ρ = −0.596, P = 5.4*10^-9^), fitness responsivity (**Supplementary Fig. S6f**; ρ = −0.413, P = 1.4*10^-4^), and expression conservation across species as measured by RNA-seq (*Ascomycota:* P = 0.00002, Mammals P = 0.0083, *Saccharomyces* P = 0.000251; two-sided Wilcoxon ranksum test). The archetypal space also distinguishes native regulatory sequences by their associated expression level (**Fig. 4g**), with intermediate expression more likely to be near the plastic archetype (A_plastic_) and depleted near the robustness cleft (**Fig. 4g**). This depletion is unlikely to result from a saturation artifact of our reporter construct; our ratiometric sorting strategy allowed us to detect saturation, but none was observed. Instead, the robustness cleft could reflect sequences at the stable extremes of one or more activation steps of gene expression (e.g. near 100% or 0% nucleosome occupied), while the plastic archetype could reflect instability around the inflection points.

Finally, we studied how natural yeast sequences explored evolutionary space. Using the 1,011 sequenced *S. cerevisiae* isolates(*67*), we placed the evolvability vectors for each set of orthologous promoters in the archetypal latent space. When a gene’s promoter from one strain is near the plastic archetype, its orthologs in the other strains tended to broadly distribute in the archetypal space (**Supplementary Fig. S6g**), but avoid the robustness cleft (*e.g*., the *DBP7* promoter from strain S288C; **Fig. 4h**). Conversely, when a promoter is near the robustness cleft (*e.g*., the *UTH1* promoter from S288C), so are its orthologs (**Fig. 4i, Supplementary Fig. S6g**). Notably, many of the native sequences in *S. cerevisiae* are near the robustness cleft (**Fig. 4j**).

In summary, the evolvability vector, which can be computed using our model directly for any sequence (without any population genetics data), encodes information about the sequence’s evolutionary history and evolvability.

## Discussion

Here, we presented a framework for addressing fundamental questions in the evolution and evolvability of *cis*-regulatory sequences(*6, 44*). The use of large scale random sequence libraries(*47*) and sensitive reporter assays(*5, 6, 80–84*) allowed us to measure the expression driven by a large number of sequences without inherent bias towards naturally occurring sequences(*19*). Using advanced deep learning approaches(*54*) and cutting edge computing hardware(*85*) for training and inference (**Methods**), we built a model from these empirical measurements that captures the complexity of *cis*-regulation and generalizes accurately in the sequence space. Our model is useful for gene-expression engineering, and can be used as an ‘oracle’ when developing and evaluating algorithms for model-guided biological sequence design(*35–39*). Importantly, we demonstrate how to use the model’s predictive power to tackle key questions in the study of fitness landscapes for understanding the genotype-phenotype-fitness relationship(*18, 19, 34*), gene expression variation across strains and species(*6*), mutational robustness(*86*), and evolvability(*44, 75*).

It has previously been suggested that evolution favors more complex regulatory solutions, with multiple weak binding sites rather than a single strong site, because complex solutions are more likely to be sampled during evolution(*87*). We showed that if stabilizing selection is acting only on gene expression, extremes of regulatory complexity gradually move towards the intermediate levels of complexity, closer to the distribution of complexity observed in native or random sequences (**Fig. 2d**). The similarity in the regulatory complexity distributions of native regulatory sequences and random sequences supports the model where most evolved regulatory sequences sample potential constraint-satisfying solutions in proportion to their frequency in the sequence space.

Recent work proposes(*88*) that adaptation to new environments can be facilitated by DNA mutations that destroy or create TF binding sites and thus cause gene mis-regulation due to regulatory crosstalk, when a TF binds the regulatory region of a gene it does not normally regulate. We found that while most sequences have one or more mutations available that are predicted to dramatically alter expression in a single environment (**Fig. 2f,g**), few mutations are available in any sequence that will satisfy competing expression objectives (**Fig. 2h**). This suggests that it would be difficult for a single promoter sequence to encode the tissue-specific expression constraints of a complex organism (where different cell/tissue types are different environments). One potential solution is to encode the regulatory activities of each gene with multiple regulatory sequences, such as the distal transcriptional enhancers that regulate cell type-specific expression in higher eukaryotes(*89*).

The d_N_/d_S_ ratio has been used extensively to characterize the evolutionary rates of protein coding genes(*90*), and we developed an analogous(*6, 64*) coefficient, the ECC, for detecting evidence of selection on gene expression from natural variation within regulatory sequences of a species. In principle, the ECC can be calculated across orthologous regulatory sequences from many different species (as opposed to individuals within a species, as we did here), but we advise caution if doing so. The ECC assumes that the function relating sequence to gene expression is the same across the orthologous sequences being compared. Since regulatory sequences evolve much faster than the regulators themselves(*10*), this assumption is likely a reasonable approximation within a species, but as evolutionary distances increase, regulators will diverge, gradually eroding this assumption. An alternative is to use gene orthology to infer the extent of expression conservation in one species using ECCs calculated in another species (**Fig. 3d**). However, such relations would extend only to well-mapped orthologs.

Complementing the ECC, which requires multiple orthologs of the regulatory region, mutational robustness as calculated with our model is predictive of selective pressures on individual sequences (**Fig. 3e,f**). While we find that strong constraint on the function of regulatory sequences can shape them to be robust to future mutations, we consider it unlikely that robustness itself is the selected trait, since increased robustness to future mutations is likely to be of little marginal benefit(*86*). Instead, this may reflect a secondary benefit of having evolved decreased expression noise(*91, 92*), or another as-yet-unknown mechanism. It may also reflect the fact that some ancestral sequences may be similar in sequence to the mutational neighbors of extant sequences, and, if selective constraints on gene expression have remained stable, these ancestral sequences likely have similar expression levels to the extant sequences.

Our approach for relating sequences using model-derived evolvability vectors allows us to study the evolutionary properties of the sequence space. Overall, we find that sequences span an evolvability spectrum from robust sequences, where few mutations alter expression appreciably and natural genetic variation tends to preserve expression, to plastic, where most mutations alter expression and natural genetic variation produced great expression diversity (**Fig. 4c,g-j**). It also helps visualize(*93*) fitness landscapes(*18*) (**Fig. 4d**, **Supplementary Fig. S7**) and future work can further improve our understanding of their global shape, dimensionality and topography(*18, 19*).

While our sequence-to-expression model produces exceptionally accurate predictions and the evolutionary insights we gained from our framework were supported by multiple lines of evidence, its direct application is currently limited by regulatory region, environment, and species. Furthermore, while we explored the interplay of competing selective pressures in two environments, most organisms are exposed to far more than two environments. In particular, for multicellular organisms, selection acts simultaneously on expression levels in many different cell types. As similar models of gene regulation are created for other species, environments, and additional regulatory regions (*e.g*. enhancers), we anticipate that the framework we presented here will continue to provide insights into *cis*-regulatory evolution.

## Acknowledgements

We thank Google TensorFlow Research Cloud for generous support in allowing us access to their Tensor Processing Units (TPUs), Broad Genomics Platform for sequencing the RNA-seq libraries, Leslie Gaffney for help with figure preparation, Jan-Christian Hütter for advice on computing the fitness responsivity, Jenna Pfiffner-Borges for yeast experiments for RNA-seq, Ruby Yu, Byron Lee and Nima Jaberi for feedback on the manuscript, Christian L. Ebbesen for the bioRxiv word template (https://github.com/chrelli/bioRxiv-word-template). and members of the Regev lab for discussions. EDV was supported by the MIT Presidential Fellowship. CGD was supported by a Canadian Institutes for Health Research Fellowship and the NIH (K99-HG009920-01). Work was supported by the Klarman Cell Observatory and HHMI. AR was an Investigator of the Howard Hughes Medical Institute.

## Conflict of Interest statement

AR is a co-founder and equity holder of Celsius Therapeutics, an equity holder in Immunitas, and until July 31, 2020 was an SAB member of ThermoFisher Scientific, Syros Pharmaceuticals, Neogene Therapeutics and Asimov. From August 1, 2020, AR is an employee of Genentech.

## Data and Code Availability

Data generated for this study are available on NCBI’s Gene Expression Omnibus, accession numbers GSE163045 and GSE163866. All models and processed data are available on Zenodo at https://zenodo.org/record/4436477 and code on GitHub at https://github.com/1edv/evolution.

## Methods

### Experimental measurement of sequence-expression pairs using a Sort-seq strategy

We experimentally measured expression using the GPRA Sort-seq(*94–97*) strategy we previously described(*47*). Briefly, for each set of expression measurements mentioned, random or designed single stranded oligonucleotides were ordered from IDT (**Supplementary Table 3**), cloned into a library as previously described(*47*) and transformed into yeast (strain Y8205 for the training dataset of random sequences, and strain *S288C::ura3* for all the rest of the sequences measured). Yeast were grown in continuous log phase, diluting as necessary to maintain an OD between 0.05 and 0.6 for 8-10 generations up until the time of harvest. Cells were harvested, washed once in ice cold PBS, and kept on ice in PBS until sorting. Cells were sorted into 18 uniformly-sized expression bins covering the majority of the expression distribution. Post sort, cells were re-grown in SD-Ura until saturation, plasmids isolated, and sequencing libraries created with a 150 cycle NextSeq kit. For libraries with random 80 bp sequences, sequences were consolidated as previously described(*47*). Reads from other (defined, non-random; synthesized by Twist Biosciences) libraries were aligned to the pre-defined sequences using Bowtie2(*98*), including only reads that perfectly matched a designed sequence. For each sequence, the expression level was the average of the expression bins in which it was observed, weighted by the number of times it was observed in each bin. These expression measurements were carried out separately in defined media lacking uracil (SD-Ura (Sunrise Science, #1703-500)) and complex media (YPD: yeast extract, peptone, dextrose).

### Architecture of the sequence-to-expression model

We captured the relationship between promoter DNA sequence (*s*) and gene expression level (*e*) as a deep transformer neural network model with the following architecture (**Supplementary Fig. S1a**):

#### Input

The input is the sequence (*s*) represented in one-hot encoding as previously described for DNA sequences(*50–53, 99–101*). Input Shape: (110,4)

#### Convolution Block

The convolution block is constructed in the following order (**Supplementary Fig. S1b**):

- Revere Complement Aware 1D Convolution. The forward and reverse strand are operated on separately with a convolutional kernel to generate strand specific sequence- environment interaction features. Kernel Shape: (30, 4, 256).
- Batch Normalization
- Rectified Linear Unit (ReLU)
- Concatenation of Features from the forward and reverse strand
- 2D Convolution: Convolve over the combined features from both the strands to capture interactions between strands. Kernel Shape: (2, 30, 4, 256)
- Batch Normalization
- ReLU
- 1D Convolution. Kernel Shape: (30, 64, 64)
- Batch Normalization
- ReLU

#### Transformer Encoder Blocks

Two transformer encoder blocks(*54*) are constructed in the following order (**Supplementary Fig. S1c**):

- Multi-Head Attention: 8 heads, capturing relations between features from different positions of (*s*) to compute a representation for the features extracted from the convolution block from (*s*).
- Residual Connection
- Layer Normalization
- Feed Forward Layer with 8 units
- Residual connection
- Layer Normalization

#### Bidirectional LSTM layer

A bidirectional LSTM layer to capture the long-range interactions between different regions of the sequence with 8 units and 0.05 dropout probability.

#### *Fully Connected Layers* (Supplementary Fig. S1d)

Two Fully connected layers with 64 Hidden Units, each consisting of ReLU and Dropout (0.05 dropout probability).

#### Output

Linear Combination of 64 features extracted as a result of all the previous operations on the sequence (*s*) to generate the predicted expression (*e*).

### Training of the sequence-to-expression model

For training, we used 20,616,659 random sequences for the defined medium and 30,722,376 random sequences for the complex medium (each to train a separate model), along with their experimentally measured expression as described above. Model architecture was written in TensorFlow(*102*) 1.14 using Python 3.6.7 with multiple open source libraries (citations, where relevant, are included in code for them). A minibatch size of 1,024 was used for training and a mean squared error loss was optimized using a RMSProp optimizer(*103*) with a learning rate of 0.001. Training was carried out on a Google Cloud Tensor Processing Unit (TPU)(*85*) v3-8. Evaluation was carried out on 4 Tesla M60 GPUs. The model architecture visualization was generated using Netron 4.5.1. All processed data and models are publicly available on Zenodo at https://zenodo.org/record/4436477 and all code is available on GitHub at https://github.com/1edv/evolution. These TPU-compatible models (for both media) were used for computing the predicted expression corresponding to a sequence throughout the manuscript unless explicitly stated otherwise in the **Methods** sections below, in which case a simpler version of the model architecture was used (which could be trained on GPUs rather than TPUs).

### Architecture of the GPU-based sequence-to-expression model

An initial model trained on GPUs (“GPU model”) used for some of the initial sequence design and evolutionary simulation sections as indicated below. This model is highly similar to that described above and used throughout most of the paper, except that it was trained using GPUs (Tesla M60s) rather than TPUs. The model did not have transformer blocks or bidirectional LSTM layers, which we incorporated into the TPU model, which required access to TPUs.

#### Input

The input is the sequence (*s*) represented in one-hot encoding as before. Input Shape: (1,110,4)

#### Convolution Block

- For the forward and reverse strand, separately,

- Strand-specific convolution layer 1. Kernel Shape: (1,30,4, 256)
- Strand-specific convolution layer 2. Kernel Shape: (30,1, 256, 256)
- Concatenation of features from the forward and reverse strand
- Convolution layer 3. Kernel Shape: (30, 1, 512, 256)
- Convolution layer 4. Kernel Shape: (30, 1, 256, 256)
- A bias term and a ReLU activation was added to each convolution layer in this block.

#### Fully Connected Layers

- Fully connected layer 1. Kernel Shape: (110*256, 256).
- Fully connected layer 2. Kernel Shape: (256, 256)
- A bias term and a ReLU activation were added to each layer in this block.

#### Output

Linear Combination of the 256 features extracted as a result of all the previous operations on the sequence (*s*) to generate the predicted expression (*e*).

Every layer was *L2* regularized with a 0.0001 weight and had a dropout probability of 0.2. A mini-batch size of 1,024 was used for training and a mean squared error loss was optimized using the Adam optimizer with an initial learning rate of 0.005. The GPU model was trained on the same data as the TPU model. Training and evaluation were carried out on 4 Tesla M60 GPUs.

### Gene expression engineering using a genetic algorithm for sequence design

To predict new sequences with desired expression we implemented a genetic algorithm (GA) with the distributed evolutionary algorithms in python (DEAP) package(*104*). The mutation probability and the two-point crossover probability were set to 0.1 and the selection tournament size was 3. The initial population size was 100,000 and the GA was run for 10 generations. The GPU model was used as the basis for the objective function for GA, which was maximized for high expression and minimized for low expression (maximizing negative predicted expression). The top 500 sequences were synthesized (by IDT) and expression was measured experimentally using our reporter assay, as described above.

### Characterizing random genetic drift

To simulate neutral mutational drift (**Fig. 2a**), we started with a set of 5,720 random sequences, in generation 0. For each sequence in this starting set, we picked a new single sequence from its 3*L* mutational neighborhood (the set of all sequences at a Hamming distance of 1 from a sequence of length *L*) randomly and calculated the difference in expression between the new sequence and the starting sequence using the model. This was done for each starting sequence to get generation 1. Each subsequent generation *n*, was produced by picking a single sequence randomly from the *3L* mutational neighborhood of each sequence in the preceding generation *n*-1. The simulation was carried out for 40 generations.

For experimental validation, we synthesized 1,000 random starting sequences, and introduced between one to three random mutations to these sequences. The expression levels of starting and mutated sequences were measured in both complex and defined media experimentally using our reporter assay. For 990 of these 1,000 starting sequences, we were able to make experimental measurements for all three mutational distances. Additionally, we introduced 20 (median) separate single mutations each to 196 native sequences, synthesized and measured their expression similarly for both of these media; these were also included in the boxes for one mutational step in **Fig. 2c** and **Supplementary Fig. S2c**.

### Characterizing the regulatory complexity of a sequence

To estimate the regulatory complexity of a sequence, we calculated the Gini coefficient of the regulatory interaction strengths for each TF. We first trained a new biochemical model with our defined media data to complement the existing one trained on complex media, using our published model architecture of TF binding and position-aware activity(*47*) and the training procedure previously described(*47*). We then individually calculated the regulatory interaction strength for each regulator by setting the concentration parameter for that TF (individually) to 0 in the learned model, and used the model to quantify the resulting change in expression, as previously described(*47*). The resulting vector of interaction strengths was used to calculate a Gini coefficient for each sequence, separately for the complex and defined media models. Regulatory complexity for a sequence is then 1-Gini. As starting points for our trajectories, we selected 200 native promoter sequences (from −160 to −80, relative to the TSS) with relatively high regulatory complexity and 200 with relatively low regulatory complexity, spanning the range of predicted expression levels, as starting points for our trajectories.

Trajectories for stabilizing selection on regulatory complexity extremes were designed using the GPU model. Here, we required all sequences to maintain a predicted expression level within 0.5 of the original expression levels at all steps along the trajectory. In order to ensure that expression was unchanged, we measured expression level experimentally for sequences along a trajectory at growing mutational steps from the initial sequence (2, 4, 8, 16, 32 mutations), as before, and excluded any trajectories where one or more of these points were missing measurements. Finally, we restricted analysis to only those trajectories for which the measured expression at no point differed from the starting measured expression level by more than 1. This resulted in a final set of 47 trajectories starting with high regulatory complexity, and 64 trajectories starting with low regulatory complexity.

### Characterizing directional trajectories under SSWM

To simulate trajectories under a Strong Selection-Weak Mutation (SSWM) regime, we started with the set of all native yeast sequences (defined as the subset from −160 to −80 relative to the TSS for all the genes in the yeast reference genome for which we had a good TSS estimate (Supplementary Table 3 in (*47*)) as the starting generation 0. For each sequence in this starting generation, we picked the sequence from its 3*L* mutational neighborhood that had the maximal (or separately, minimal) predicted expression using our model to get generation 1. Each subsequent generation *n* was produced by picking for each sequence in generation *n*-1 the sequence from its 3*L* mutational neighborhood with the maximal (or separately, minimal) expression. The simulation was carried out for 10 rounds.

For experimental validation, we synthesized a subset of sequences from several generations along simulated mutational trajectories using the GPU model for defined (6,304 sequences from 637 trajectories, 591 of which had every sequence along the trajectory successfully measured) and complex media (10,322 sequences from 877 trajectories, 805 of which had every sequence along the trajectory successfully measured) and measured their expression in the corresponding media experimentally using our reporter assay.

### Measuring the *URA3* expression-to-fitness relationship

We studied two complementary environments with opposite selective pressures on the expression of *URA3* (encoding an enzyme responsible for uracil synthesis): defined media, where organismal fitness increases with gene expression (up to saturation) and complex media + 5-FOA, where fitness decreases with Ura3 expression.

We used the GPU models trained on defined and complex media to choose a set of 11 sequences that span a broad range of predicted expression levels in the two media when cloned into a YFP expression vector(*47*). We experimentally estimated the relationship between expression of *URA3* and organismal fitness in yeast, from these 11 sequences, by cloning promoter sequence in front of YFP to measure expression level and in front of *URA3* to measure fitness. Unless otherwise noted, yeast were grown at 30°C, in an orbital shaker incubator at 225 RPM. Each vector was transformed into yeast (S288C::*urα3*), and three independent transformants were selected per vector to serve as biological replicates. For measuring expression, yeast were grown overnight in either YPD+NAT (yeast extract, peptone, dextrose, with 75μg/ml nourseothricin) or SD-Ura (synthetic defined media, lacking uracil; Sunrise Science 1703-500), and then re-inoculated in the morning and allowed to grow for 6 hours prior to measuring expression by flow cytometry for each replicate as the log ratio of YFP to the constant background RFP, including only cells obtaining the top 50% of RFP expression. Fitness was obtained by measuring the growth rate of each yeast strain in either SD-Ura or YPD+NAT+5-FOA (0.25 mg/ml 5-FOA). Yeast were grown continuously in triplicate in log phase, with linear shaking at 30°C in a Synergy H1 plate reader (Biotek), by diluting each well to maintain OD<0.7, with OD measured at 15 minute intervals. Growth rate was defined for each replicate as the median of the instantaneous smoothed growth rates over 5 measurements in log phase, considering only time points where 0.05<OD<0.5. Each promoter’s expression and growth rate were summarized as the mean of the three replicates.

### Characterizing trajectories under conflicting expression objectives in different environments

To simulate sequence evolution in two complementary environments with opposite selective pressures (defined media and complex media), we started with the set of all native yeast sequences as the starting generation 0, and defined the objective function as the difference in predicted expression between defined and complex media using the models trained in the respective media. In one experiment, we maximized this difference (defined minus complex), and the other we minimized it (maximizing complex minus defined). For each sequence in generation 0, we picked the sequence from its 3*L* mutational neighborhood that had the maximum (or separately, minimum) value for the objective function as generation 1 using the model. Each subsequent generation *n* was produced by picking for each sequence in generation *n*-1 the sequence from its 3*L* mutational neighborhood with the maximum (or separately, minimum) value for the objective function, to a total of 10 generations.

We *de novo* identified motifs that were enriched in the sequences of generation 10 compared to the starting sequences using DREME(*105*), and searched each of the top 5 consensus motifs in the YeTFaSCo database(*106*), reporting the closest match, or one of multiple similar matches.

### Characterizing Finding orthologous promoters in the 1,011 *S. cerevisiae* genomes dataset

To identify orthologs of S288C promoters in the whole genome sequences of the 1,011 yeast strains(*67*), we used BLAT(*107*) to identify regions of ≥80% identity with each −160 to −80 region (relative to the TSS) annotated in the reference S288C genome sequence (R64)(*108*). We excluded on a gene-by-gene basis any strains with more than one such match, where the match contained insertions or deletions, or had incomplete matches. Genes with more than 1.2 matches with ≥80% identity per genome, on average, were excluded altogether.

### Computing the expression conservation coefficient (ECC)

To calculate the ECC, for each yeast gene promoter, we used the model to predict an expression value for each orthologous promoter in the 1,011 yeast genomes (above), defining an expression distribution with a standard deviation *σ_B_*. We also generated, from each gene’s consensus promoter sequence (defined as the most abundant base at each position across the strains), a set of sequences with random mutations, such that the number of sequences at each Hamming distance from the consensus promoter sequence was the same for the natural and simulated sets. We used the same model to predict the expression of the simulated sequences, and calculate its standard deviation *σ_C_*. The nominal ECC is *log(σ_C_/σ_B_*). Because the variance on simulated sequences is better estimated than in natural orthologs (whose sequences may be more constrained), we subtract a constant correction factor calculated by creating a second simulated set of randomly mutated sequences whose diversity is limited to the same extent as in the natural set, by creating only one random mutation for every unique sequence in the set of native orthologs. We then predict expression for this second set, and use this standard deviation (*σ_C_*) to calculate a null ECC for each gene (*log(σ_C_/σ_C_))*; he median of these null ECCs over all the genes is used as the constant correction factor 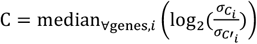

The corrected ECC for gene*g* is then:

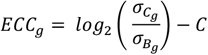

The computed ECC values for all yeast genes, available in **Supplementary Table 1**, were used to identify cases or presumed stabilizing selection (selection favoring a fixed non-extreme value of a trait), diversifying (disruptive) selection (selection favoring more than one extreme values of a trait; as opposed to a single fixed intermediate value), and directional (positive) selection (selection favoring a single extreme value of a trait over all other possible values of the trait). We re-computed the ECC values for all yeast genes using the S288C reference sequences instead of the consensus sequence for the promoters of each gene and got very similar results.

### Inferring expression conservation across *Saccharomyces* species using RNA-seq data and comparing with ECC values

Published RPKM values for orthologs of *S. cerevisiae* genes in closely related *Saccharomyces* species(*71*) were obtained from the Gene Expression Omnibus (GEO) (accession GSE83120). Only genes for which expression was quantified in all species were used in subsequent analysis. RPKM values were log_2_ scaled after adding a pseudo count of 2, and the variance in expression of each gene across the species was calculated. We ranked genes by their gene expression variance, and took the 2% of genes with the lowest variance as those considered to have conserved gene expression levels (‘expression conserved’), while the 2% with the highest variance were considered ‘expression not-conserved’. T o compare to ECC values, we estimated the p-value of a two-sided Wilcoxon rank-sum test (implemented using the *scipy.stats.ranksums* SciPy(*109*) function) comparing the ECC values for genes in the ‘expression conserved’ and ‘expression not-conserved’ categories. To control for the dependence between expression mean and variance, we also repeated the analysis using the coefficient of variation (P = 1.05*10^-4^) and the coefficient of dispersion (P = 2.42*10^-4^) instead of variance and obtained similar results.

### Experimental protocol for RNA-seq measurements from 11 Ascomycota species

We performed RNA-seq on the following 11 Ascomycota yeast species: *Saccharomyces cerevisiae, Saccharomyces bayanus, Naumovozyma (Saccharomyces) castellii, Candida glabrata, Kluyveromyces lactis, Kluyveromyces waltii, Candida albicans, Yarrowia lipolytica, Schizosaccharomyces japonicus, Schizosaccharomyces octosporus*, and *Schizosaccharomyces pombe*. Each of the 11 species was grown in BMW medium, chosen to minimize cross-species growth differences, as previously described(*110*). *N. castellii* was grown at 25°C while the rest of the species were grown at 30°C. RNeasy Midi or Mini Kits (Qiagen, Valencia, CA) were used to isolate total RNA from log-phase cells by mechanical lysis using the manufacturer instructions as previously described(*110*). dUTP strand-specific RNA-seq libraries were constructed as previously described(*111*) with the following modifications. (**1**) The polyA^+^-selected RNA was fragmented in a 40 μl reaction containing 1x Fragmentation Buffer (Affymetrix) by heating at 80°C for 4 minutes followed by cleanup via ethanol precipitation for all libraries (except *Y. lipolytica, S. pombe, S. japonicus*, and *S. octosporus;* for these species, the conditions described previously were used(*111*)), followed by cleanup via 1.8x RNAClean XP beads (Beckman Coulter Genomics). (**2**) For *C. glabrata, K. lactis, S. bayanus, S. pombe, S. japonicus*, and *S. octosporus* libraries, the adapter ligation was performed overnight at 16°C. For the rest, this was done at 16°C for 2 hours as described previously(*111*). (**3**) Normalization was carried out based on the cDNA input and pooling of selected Illumina barcoded-adaptor-ligated cDNA products followed by gel size selection occurred as follows: range of 275 to 575 bp for pooled *C. albicans, K. waltii*, and *N. castellii* libraries, and 375 to 575 bp for *C. glabrata, K. lactis*, and *S. bayanus* libraries. For the other libraries, no pooling was performed before gel size-selection – range of 310 to 510 bp for *Y. lipolytica* and 350 to 550 bp for *S. pombe, S. japonicus*, and *S. octosporus*. (**4**) The final PCR product was purified by 1.8x AMPure XP beads (Beckman Coulter Genomics) followed by a second gel size-selection for the range of 300 to 575 bp for *C. albicans, K. waltii*, and *S. castellii* libraries, but no second gel size-selection was performed for the other libraries. The pooled final library was sequenced on one to four lanes of HiSeq2000 (Illumina) with 68 base (*Y. lipolytica* had 76 base) paired-end reads and 8 base index reads.

### Transcript assembly, mapping and expression calculation for the 11 Ascomycota species RNA-seq

For each of the 11 Ascomycota yeast species above, reads were assembled using Trinity(*112*)(version ‘trinityrnaseq_r2012-05-18’) and the assembled transcripts were mapped onto the assemblies to the respective genomes using GMAP(*113*). The Jaccard coefficient was used to join adjacent assemblies given enough connecting reads (using the Trinity default of 0.35 for the Jaccard cutoff). Finally, upon mapping all assembled transcripts, the Jaccard coefficient was used to clip assemblies which did not have enough support over a certain region. For each of the species, assembled transcripts were mapped to the genome sequence(*114*) using BLAT(*107*). Estimated expression values were calculated for each transcript using RSEM(*115*) (defined in RSEM as the estimate of the number of fragments that are derived from a given isoform or gene, or the expectation of the number of alignable and unfiltered fragments that are derived from an isoform or gene given the maximum likelihood abundances). Only reads mapping to the sense mRNA strand were considered. Orthology between genes in different species was used as previously described(*114*).

### Inferring expression conservation across Ascomycota species using our RNA-seq data and comparing with ECC values

Estimated expression values from the 11 Ascomycota species RNA-seq data were used after removing all genes with NA values in expression for more than three species. Estimated expression values were *log*_2_ scaled after adding a pseudo count of 1, and the variance in expression for each gene across the species was calculated. Genes were ordered by their variance in expression across the reported fungal species. Here, the 10% of genes with the lowest expression variance were considered to have ‘conserved’ expression, and the 10% with highest expression variance were considered to have expression ‘not conserved’. To compare to ECC values, we estimated the p-value of a two-sided Wilcoxon rank-sum test (implemented using the *scipy.stats.ranksums* SciPy(*109*) function) comparing the ECC values for genes in the ‘conserved’ and ‘not conserved’ categories. We obtained similar results when we repeated the analysis using the coefficient of variation (P = 4.22*10^-5^) and the coefficient of dispersion (P = 8.05*10^-5^) instead of variance.

### Inferring expression conservation across Mammalian species using RNA-seq data and comparing with ECC values

Ensembl Biomart(*116*) was used to find one to one or one to many orthologs of *S. cerevisiae* genes in humans (of ‘Human homology type’ either ‘ortholog_one2one’ or ‘ortholog_one2many’; all ‘many2many’ orthologs were excluded). A percent identity >50% (‘%id. query gene identical to target Human gene’) was also required for an ortholog pair to be used in the subsequent analysis. For the retained human orthologs of yeast genes, we directly used the previously reported ‘evolutionary variance’ values across mammalian species from the original publication(*72*) (based on an Ornstein Uhlenbeck (OU model)(*72*)). Here, the 25% of genes with the lowest ‘evolutionary variance’ were considered to have conserved expression and the top 25% were considered to be not conserved (the same thresholds used in the original study(*72*)). This was done separately for each profiled tissue (brain, heart, kidney, liver, lung and skeletal muscle). We verified that were no genes with conflicting expression conservation classes among tissues. Subsequently, a human ortholog for a yeast gene was considered to have conserved (or non-conserved) expression if it was found to have conserved (or non-conserved) expression in at least one of the profiled tissues. To compare to ECC values, we estimated the p-value of a two-sided Wilcoxon rank-sum test (implemented using the *scipy.stats.ranksums* SciPy(*109*) function) comparing the ECC values for genes in the “conserved” and “not conserved” categories.

### Quantifying sequence dissimilarity using mean Hamming distance

For each group of orthologous yeast gene promoters (with ungapped alignments), we calculated the mean of Hamming distances between each pair of orthologous promoters across the 1,011 isolates.

### Fitness responsivity

Published expression-to-fitness curves in glucose media for each of 80 genes were obtained from the Supplementary Data of the original publication(*22*). For each of these curves, the total variation (**Supplementary Fig. S5**) was calculated by partitioning the expression range into 36 regular intervals (as reported in the ‘impulse fit’ of the expression-to-fitness curves in the original publication(*22*)) and summing the absolute difference in fitness at the endpoints of each partition as follows correction factor 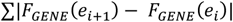, for each gene’s expression-to-fitness function, *F_GENE_*^(e)^.

### Mutational robustness

For every sequence, mutational robustness was defined as the fraction of sequences in its 3*L* mutational neighborhood that altered the expression by an amount less than *ε*, where *ε* is set at two times the standard deviation of expression variance across all genes with an ECC >0 (here, ε = 0.1616; ECC calculated using the 1,011 *S. cerevisiae* genomes, **Supplementary Fig. S4d**). Using different values for ε yielded very similar results.

### The evolvability vector

To compute an evolvability vector for a sequence *S*_0_, for each sequence *S_i_* in the 3*L* mutational neighborhood of *S*_0_, we calculate the difference between the predicted expression of *S_i_* and that of *S*_0_: *d_i_* = *f*(*S_i_*) — *f*(*S*_0_), where *f*(*S*) represents the predicted expression of the model. We define the evolvability vectors as the vector D ({d_1_, *d_2_*, …, d_3L_}), sorted such that *d_i_* ≥ *d*_*i*-1_, ∀*i* (i.e. *d_i_* values are in ascending order).

### Archetypal analysis of the sequence space using evolvability vectors

The evolvability vectors for a new random sample of a million sequences were used as input to an autoencoder with an archetypal regularization constraint(*79*) on the embedding layer. The autoencoder was trained using the AANet implementation made available with the publication(*79*) with no noise added to the archetypal layer during training, a linear activation on the output layer, an equal weight of 1 on each of the loss terms (the mean squared error loss term along with the non-negativity and convexity constraints), a learning rate of 0.001, and a minibatch size of 4,096. The autoencoder accepts an evolvability vector (of length 240 for an 80bp sequence) as input to the first encoder layer, where each node in the input layer is connected to each node in the encoder layer (fully connected layer). Every layer in the autoencoder was fully connected. The encoder architecture used was [1024,512,256,128,64] where each entry corresponds to the number of nodes in the corresponding hidden layer and the decoder architecture was the encoder’s mirror image. The output layer was the same shape as input layer and each node in the last decoder layer was connected to each node in the output layer. To select the optimal number of archetypes, the autoencoder was first trained for a 1,000 minibatches separately for 1 to 9 archetypes. Following the recommended approach(*79*) for picking the optimal number of archetypes, we used an elbow plot of mean squared error on the evolvability vectors (here, using native sequences) *vs*. the number of archetypes in the autoencoder (**Supplementary Fig. S6a**).

We then trained the autoencoder from scratch with 3 archetypes, using the full training data and parameters for 250,000 batches. Since this autoencoder aims to reconstruct the original evolvability vector for each sequence by learning feature representations after passing them through an information bottleneck, we first verified its reconstruction accuracy on the set of native yeast promoter sequences (**Supplementary Fig. S6b**, Pearson’s r = 0.992). To visualize the evolvability vectors corresponding to sequences in 2 dimensions (2D), the evolvability vectors corresponding to the three archetypes were first generated by decoding their archetypal latent space coordinates ((1,0,0), (0,1,0) and (0,0,1)) through the decoder, and MDS was performed on the decoded evolvability vectors of the archetypes. Then, as previously described(*79*), the encoded evolvability vector of each new sequence was projected into the 2D MDS space by representing it as a mixture of the archetypes and interpolating them between the MDS coordinates of each archetype. For every sequence, we can now compute the following equivalent representations: (i) its evolvability vector, (ii) an archetypal triplet quantifying the similarity of its encoded (latent space) evolvability vector to the three archetypes and (iii) a two-dimensional multidimensional scaling (MDS) coordinate(*79*) for visualizing the evolvability vectors. The representation of the evolvability vector for each sequence in this archetypal space is now bounded by a simplex (whose vertices correspond to the 3 evolvability archetypes). For each native and natural yeast promoter sequence from the sequence space, we inferred the archetypal triplet and MDS coordinates using its evolvability vector with this trained autoencoder. The MDS coordinates for the archetypes and the native yeast promoter sequences were used to generate the visualizations of the sequence space shown.

### Visualizing promoter fitness landscapes

1000 random sequences were sampled and projected onto the MDS coordinate system for visualizing the sequence space described above. The expression level of each sequence was calculated using our model, and expression values were scaled so that the minimum was 0 and maximum was 1. Previously quantified expression-to-fitness relationships(*22*) to compute fitness (fraction of wildtype growth rate) by using cubic spline interpolation (implemented using the *scipy.interpolate.CubicSpline* SciPy(*109*) function) on the expression level after scaling the measured expression-to-fitness curves to have an expression range of 0 to 1. These fitness values were then used to generate the contour plots (implemented using the *matplotlib.pyplot.tricontourf* function; **Fig. 4d**, **Supplementary Fig. S7**) that visualize the fitness landscape in that gene’s promoter sequence space.

## Supplementary Tables

### Supplementary Table 1

The Expression Conservation Coefficient (ECC), mutation tolerance, evolvability vector archetypal coordinates and predicted expression corresponding to all native promoter sequences.

### Supplementary Table 2

The GO terms enriched by the ECC ranking.

### Supplementary Table 3

The list of single stranded oligonucleotides used.

---

**Supplementary Fig. S1.**
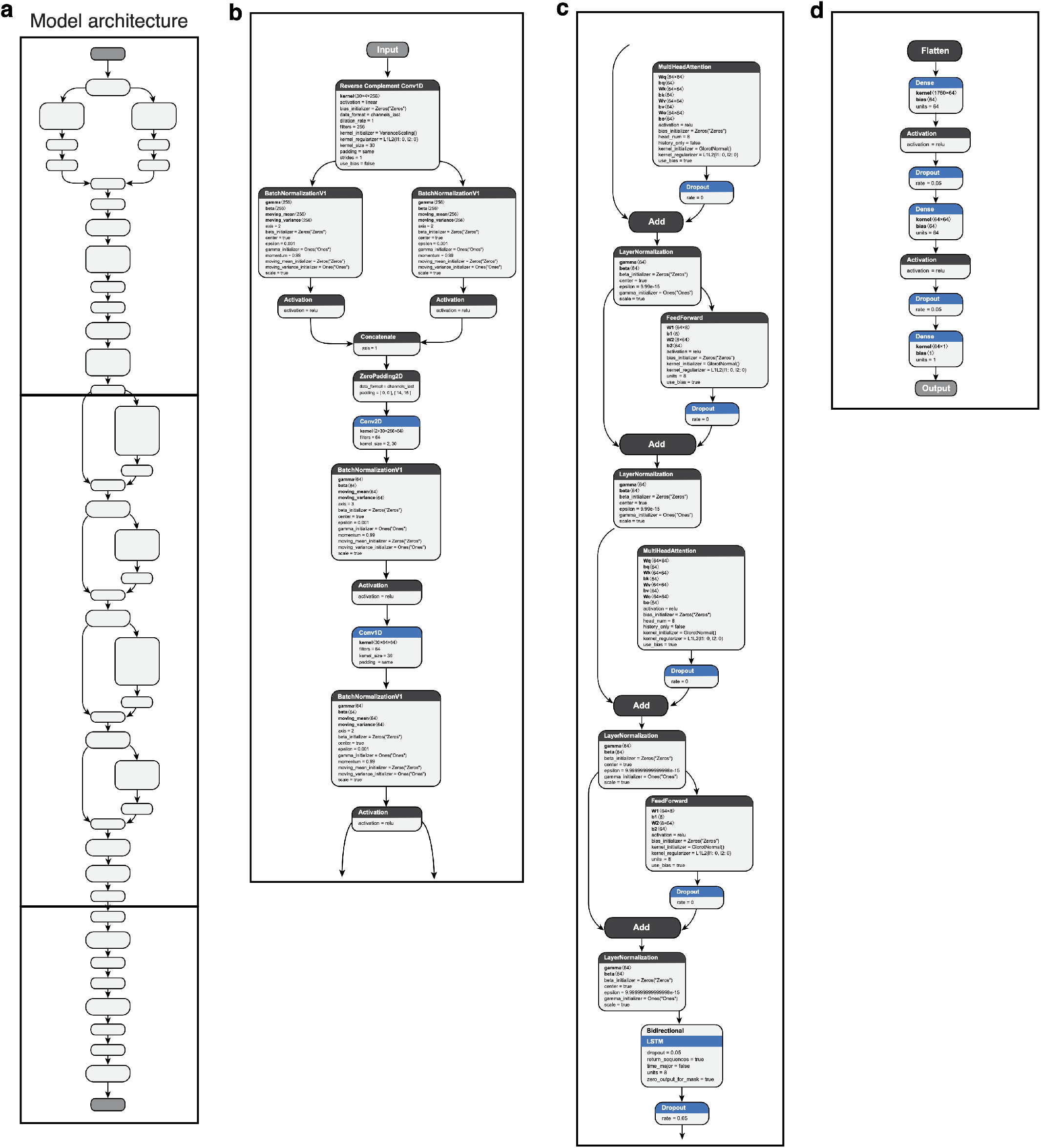
The deep transformer neural network architecture for the sequence-to-expression model. **a,** Model architecture with three blocks (horizontal lines) and multiple layers (boxes). b-d. Expanded architecture (**Methods**) for the convolutional (**b**), transformer encoder (**c**) and multi-layer perceptron (**d**) blocks.

**Supplementary Fig. S2.**
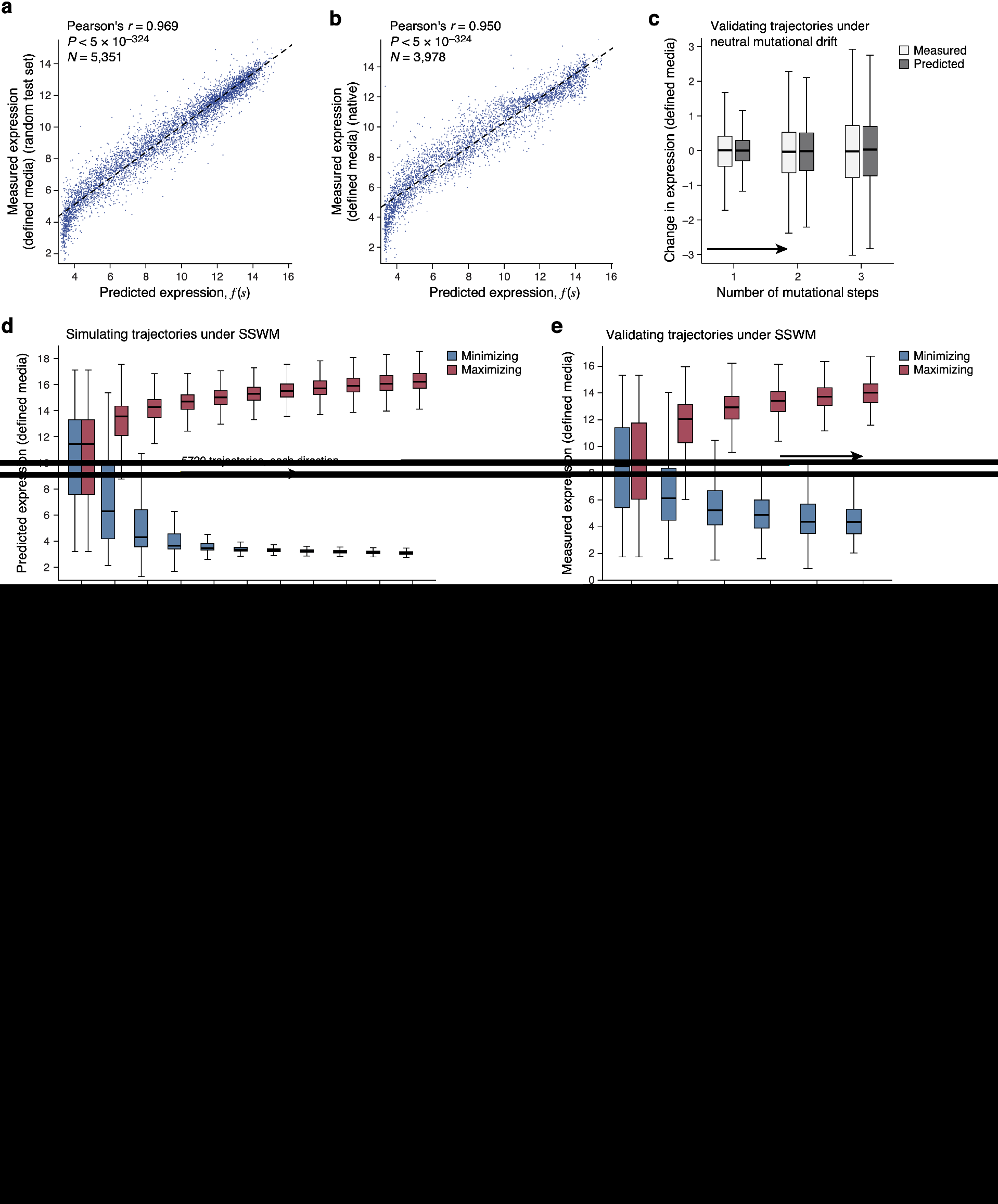
The sequence-to-expression model generalizes accurately and helps characterize sequence trajectories under different evolutionary regimes. **a,b,** Accurate prediction of expression from sequence in defined media. Predicted (*x* axis) and experimentally measured (*y* axis) expression in defined media (SD-Uracil) for **(a)** random test sequences (sampled separately from and not overlapping with the training data) and **(b)** native yeast promoter sequences. Top left: Pearson’s *r* and associated P-value. **c,** Experimental validation of trajectories from simulations of random genetic drift. Distribution of measured (light grey) and predicted (dark gray) changes in expression in the defined media (SD-Uracil) (*y* axis) for the synthesized sequences at each mutational step (*x* axis) from predicted mutational trajectories under random mutational drift. Midline: median; boxes: interquartile range; whiskers: 1.5x interquartile range. **d, e,** Simulation and validation of expression trajectories under SSWM in defined media (SD-Uracil). **d,** Distribution of predicted expression levels (*y* axis) in defined media at each evolutionary time step (*x* axis) for sequences under SSWM favoring high (red) or low (blue) expression, starting with 5,720 native promoter sequences. Midline: median; boxes: interquartile range; whiskers: 1.5x interquartile range. **e,** Experimentally measured expression distribution in defined media (*y* axis) for the synthesized sequences at each mutational step (*x* axis) from predicted mutational trajectories under SSWM, favoring high (red) or low (blue) expression. Midline: median; boxes: interquartile range; whiskers: 1.5x interquartile range. **f-m,** Experimental validation of predicted expression for sequences from the random genetic drift and SSWM simulations. Experimentally measured (*y* axis) and predicted (*x* axis) expression level (**j-m**) or expression change from the starting sequence (**f-i**) in complex (**f**,**j**,**h**,**l**) or defined (**g**,**i**,**k**,**m**) media using sequences from the random drift (**Fig. 2c** and **(c)**; **f**,**g**,**j**,**k** here) and SSWM (**Fig. 2g** and (**d,e**); **h**,**i**,**l**,**m** here) simulations. Top left: Pearson’s *r* and associated P-values.

**Supplementary Fig. S3.**
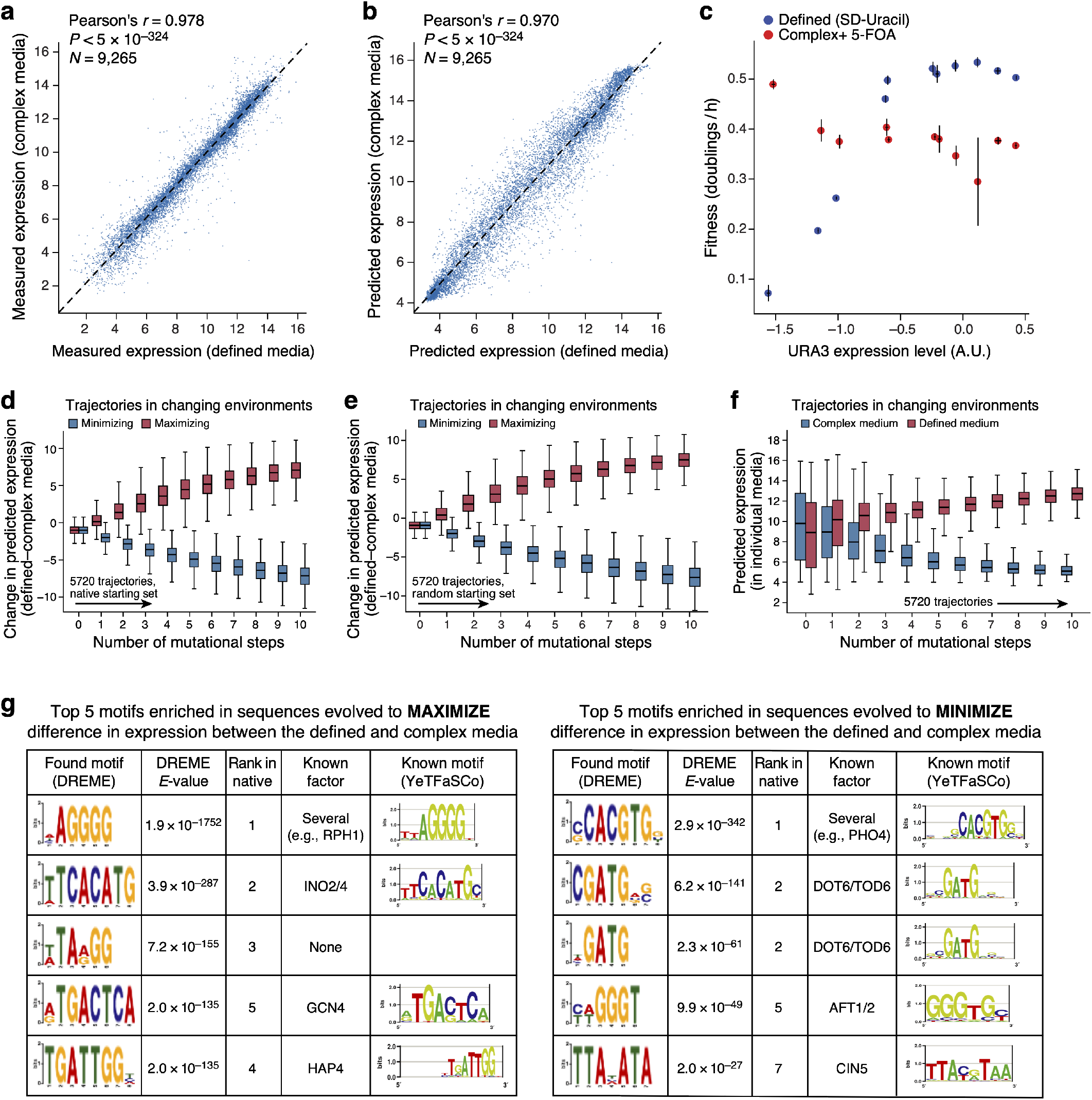
Characterization of sequence trajectories under strong competing selection pressures. **a,b,** Expression is highly correlated between defined and complex media. Measured (**a**) and predicted (**b**) expression in defined (*x* axis) and complex (*y* axis) media for random test sequences. **c,** Opposing relationships between organismal fitness and URA3 expression in two environments. Measured expression (x axis, using a YFP reporter) and fitness (*y* axis; when used as the promoter sequence for the *URA3* gene) for yeast with each of 11 promoters predicted to span a wide range of expression levels in complex media with 5-FOA (red), where higher expression of *URA3* is toxic due to *URA3*-mediated conversion of 5-FOA to 5-fluorouracil, and in defined media lacking uracil (blue), where URA3 is required for uracil synthesis. Error bars: Standard error of the mean. **d-f,** Competing expression objectives are slow to reach saturation. **d,e,** Difference in predicted expression (*y* axis) at each evolutionary time step (*x* axis) under selection to maximize (red) or minimize (blue) the difference between expression in defined and complex media, starting with either native sequences (**d,** as **Fig. 2f**) or random sequences (**e**). **f,** Distribution of predicted expression (*y* axis) in complex (blue) and defined (red) media at each evolutionary time step (*x* axis) for a starting set of 5,720 random sequences. Midline: median; boxes: interquartile range; whiskers: 1.5x interquartile range. **g,** Motifs enriched within sequences evolved for competing objectives in different environments. Top five most enriched motifs, found using DREME(*105*) (**Methods**) within sequences computationally evolved from a starting set of random sequences to either maximize (left) or minimize (right) the difference in expression between defined and complex media, along with DREME E-values, the corresponding rank of the same motif when using native sequences as a starting point, the likely cognate TF and that TF’s known motif.

**Supplementary Fig. S4.**
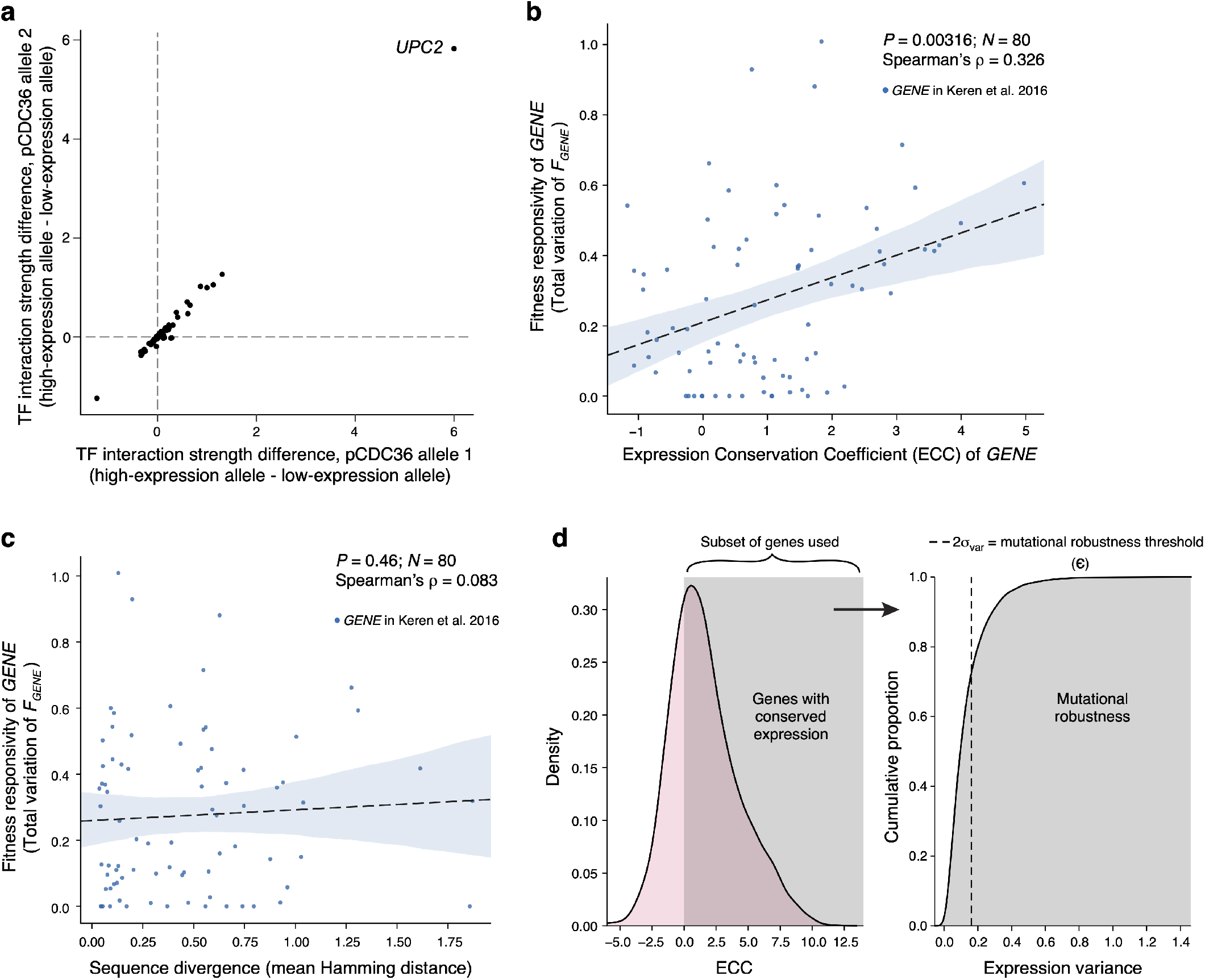
Signatures of stabilizing selection on gene expression detected from regulatory DNA across natural populations. **a,** Expression-altering alleles in the CDC36 promoter are attributed primarily to altered UPC2 binding. TF interaction strength(*47*) (expression attributable to each TF) difference between the high and low alleles (each point is a TF) for each of two low expression alleles (allele 1: *x* axis; allele 2: *y* axis). Each low-expressing allele is compared to the high-expression allele with the most similar sequence (across all promoter sequences analyzed from the 1,011 strains; *e_TF,A_high__* — *e_TF,A_low__*). **b, c,** Fitness responsivity is associated with ECC, but not with simple sequence diversity. Fitness responsivity (*y*axes) and ECC (**b,** *x*axis) or mean Hamming distance (**c,** *x*axis) for each of 80 genes (points). Top right: Spearman’s *ρ* and associated P-values. **d**, Determination of expression change threshold for defining a “tolerated mutation” to compute mutational robustness. We used all genes with an ECC consistent with stabilizing selection (ECC>0; left), calculated the variance in predicted expression across the 1011 yeast strains for each gene, and chose the tolerable mutation threshold, *ε*, as two standard deviations of the distribution of the variance (right). ~73% of genes with ECC>0 had an expression variation lower than ∈.

**Supplementary Fig. S5.**
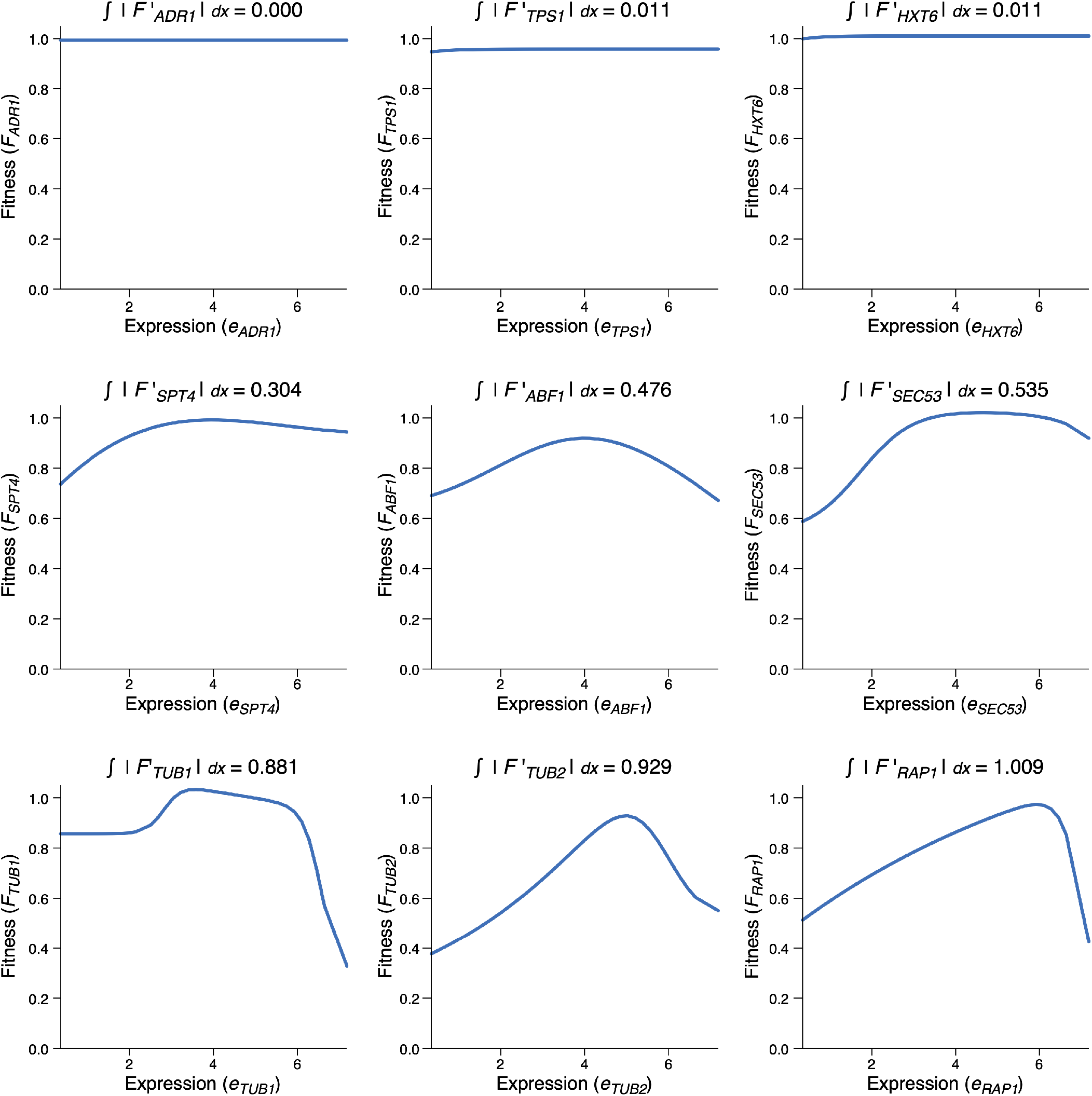
Fitness responsivity of a gene as the total variation of its expression-to-fitness relationship *F_GENE_* curves. Expression (*x* axis) and fitness (*y* axis) levels for different promoter variants for each select gene fit from experimental measurements by Keren et al(*22*). Fitness responsivity calculated as the total variation in each curve is noted above each panel.

**Supplementary Fig. S6.**
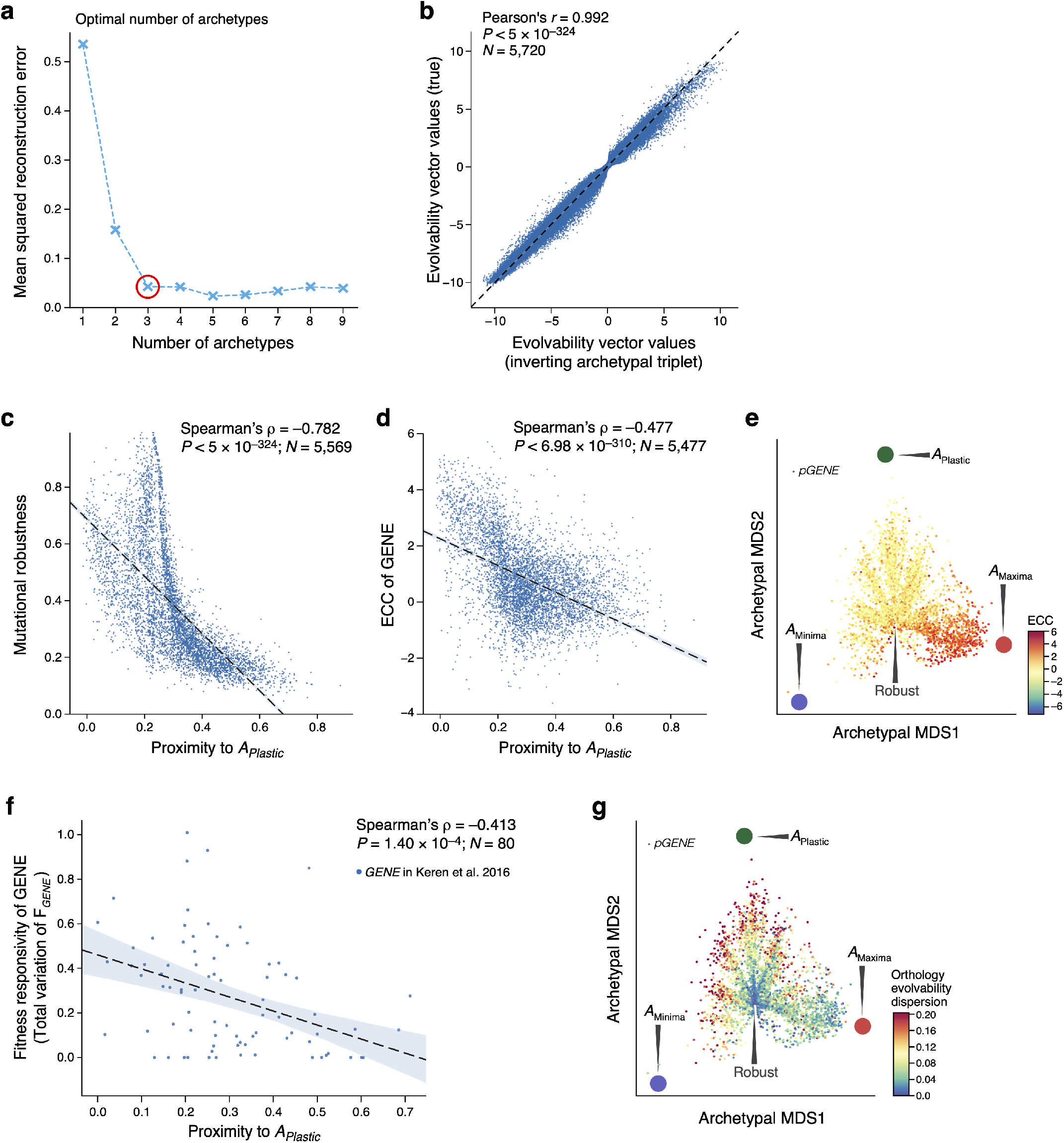
Analysis of cis-regulatory evolvability reveals sequence-encoded signatures of expression conservation from solitary sequences. **a,** Selection of optimal number of archetypes. Mean-square-reconstruction error (*y* axis) for reconstructing the evolvability vectors from the embeddings learned by the autoencoder for an increasing number of archetypes (*x* axis). Red circle: optimal number of archetypes selected as prescribed(*79*) by the “elbow method”. **b,** The archetypal embeddings learned by the autoencoder accurately capture evolvability vectors. Original (*y* axis) and reconstructed (*x* axis) expression changes (the values in the evolvability vectors) for each native sequence (none seen by the autoencoder in training). **c,d**, Sequence-encoded signatures of expression conservation. The proximity to the plastic archetype (*x* axes) and mutational robustness (*y* axis, **c**) or ECC (*y* axis, **d**) for all yeast genes. Top right: Spearman’s *ρ* and associated P-values. “L”-shape of relationship in **c** results from the robust cleft, A_Maxima_, and A_Minima_ all being distal to A_Plastic_ (left side of plot). **e,** All native (S288C reference) promoter sequences (points) projected onto the evolvability archetype space learned from random sequences; colored by their ECC. Large colored circles: evolvability archetypes. **f**, The proximity to the plastic archetype (*x* axis) and fitness responsivity (*y* axis) for the 80 genes with measured fitness responsivity. Top right: Spearman’s *ρ* and associated P-values. **g,** All native (S288C reference) promoter sequences (points) projected on the evolvability archetype space learned from random sequences; colored by their mean pairwise distance in the evolvability archetype space between all promoter alleles across the 1,011 yeast isolates for that gene (ortholog evolvability dispersion). Large colored circles: evolvability archetypes.

**Supplementary Fig. S7.**
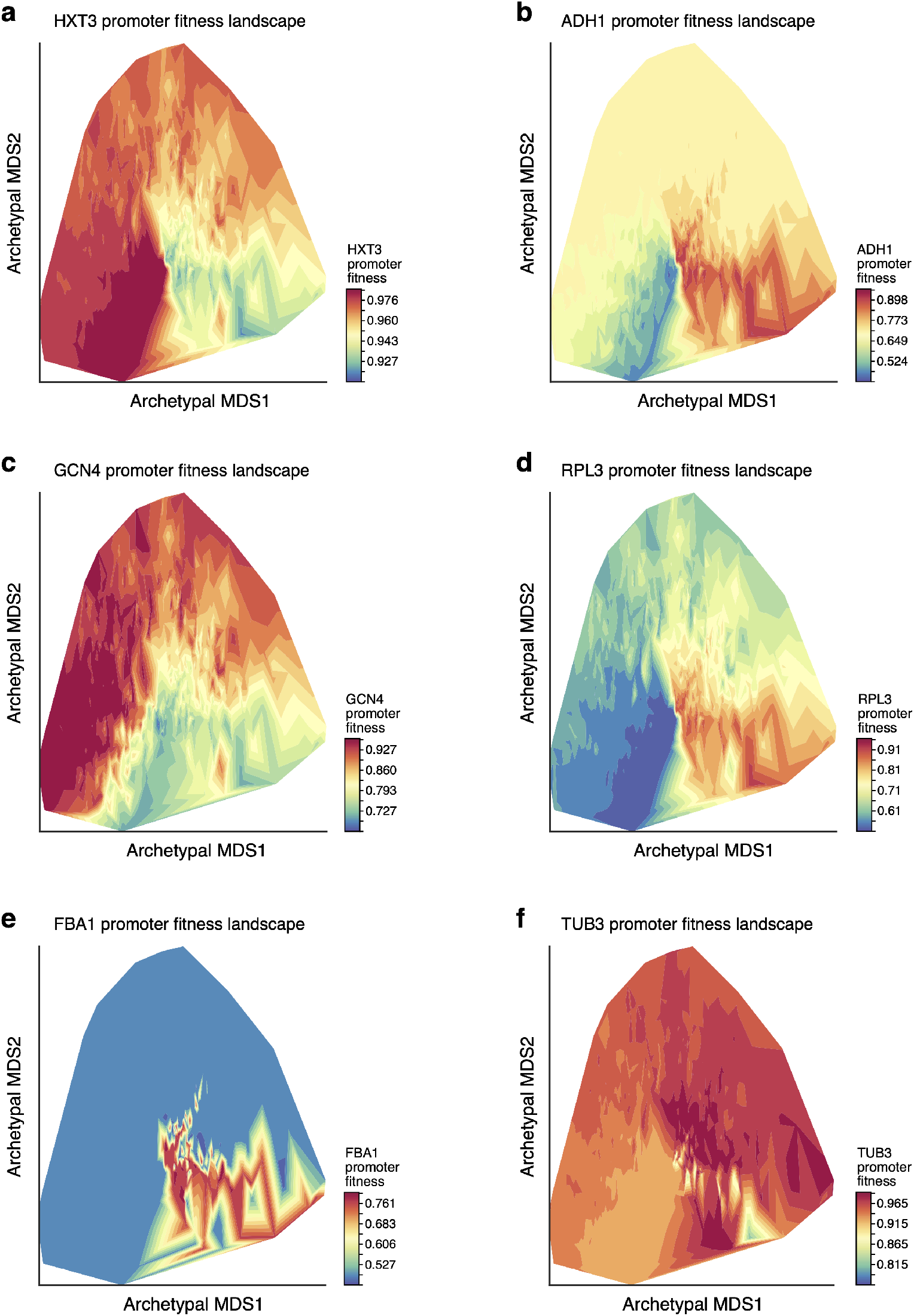
Visualizing promoter fitness landscapes in sequence space. Visualizing the fitness landscapes for the promoters of HXT3 (**a**), ADH1 (**b**), GCN4 (**c**), RPL3 (**d**), FBA1 (**e**), TUB3 (**f**). 1000 promoter sequences represented by their evolvability vectors projected onto the 2D archetypal space and colored by their associated fitness as reflected by their predicted growth rate relative to wildtype (color, **Methods**), estimated by first mapping sequences to expression with our model and then expression to fitness as measured and estimated previously(*22*).

## Notes

https://zenodo.org/record/4436477

https://github.com/1edv/evolution

### Summary of Updates

References updated based on community feedback; biorxiv logo on PDF now links to DOI; Panel titles reformatted on Supplementary Fig. S5;

